# Natural variation in NifU and NifS enhances chloroplasts compatibility for nitrogenase engineering

**DOI:** 10.64898/2026.07.09.737459

**Authors:** Manuel Ene-Ordorica, Cristina Vaca-Sanz, Natalia Makarovsky-Saavedra, Adrian O. Sanchez, Francesco Blasio, Leonardo Curatti, Elena Caro, Luis M. Rubio

**Affiliations:** Centro de Biotecnología y Genómica de Plantas, Universidad Politécnica de Madrid e Instituto Nacional de Investigación y Tecnología Agraria y Alimentaria/Consejo Superior de Investigaciones Científicas, Madrid 28223, Spain; Instituto de Investigaciones en Biodiversidad y Biotecnología, INBIOTEC-CONICET and Fundación para Investigaciones Biológicas Aplicadas, Mar del Plata, Buenos Aires, Argentina

**Keywords:** Nitrogen fixation engineering Plant biotechnology, Rice transformation Fe-S cluster pathways, Plastid protein expression Protein solubility, Rice proteome Chloroplast physiology Oxidative stress

## Abstract

Reconstitution of functional nitrogenase in plants requires the coordinated expression of the [Fe–S] cluster assembly proteins NifU and NifS. However, the extent to which these proteins interact with endogenous Fe–S metabolism and affect plant physiology remains unclear. Here, we compared NifU and NifS homologs from diverse diazotrophs to identify variants compatible with the plant chloroplast environment. Selected variants of *Azotobacter vinelandii*, *Fischerella thermalis*, and *Marinobacter lutimaris* were characterized by transient expression in *Nicotiana benthamiana* and stable transformation in rice. Plant-produced NifU was largely devoid of [Fe–S] clusters when isolated but retained strong capacity for *in vitro* [Fe–S] cluster reconstitution and apo-NifH activation in a *Ft* > *Av* >*Ml* gradient, indicating correct folding and function but limited cluster loading or stability *in vivo*. NifU and NifS expression in transgenic rice resulted in variant-dependent proteome and phenotype effects, with *A. vinelandii-*expressing lines exhibiting severe defects, *F. thermalis* lines showing intermediate phenotype, and *M. lutimaris* lines being indistinguishable from wild type. These results reveal a trade-off between the biochemical activity of NifU and NifS and their compatibility with host metabolism, which must be considered for successful nitrogenase engineering in plants.

**Highlight:** NifU/NifS homolog selection determines trade-offs between [Fe–S] cluster assembly activity and plant compatibility, identifying variants that minimize physiological disruption while supporting nitrogenase cofactor assembly in chloroplasts.

## Introduction

Rice, along with other cereal crops such as wheat and maize, provides nearly half of the calories consumed directly by humans (Erisman *et al*., 2008; Mueller *et al*., 2012). Cereal crop productivity is highly dependent on nitrogen availability in the soil, and almost entirely on synthetic nitrogen fertilizers produced by the high-energy-cost Haber–Bosch process to sustain their yields (Tilman *et al*., 2002). Unfortunately, widespread N fertilizer application contributes to eutrophication, biodiversity loss, and the accumulation of nitrous oxide, a potent greenhouse gas (Erisman *et al*., 2015; Zhang *et al*., 2015). Furthermore, fertilizer costs are an economic hurdle for smallholder farmers in some regions of the world, such as sub-Saharan Africa and Southeast Asia (Mueller *et al*., 2012). Among the different biological strategies to improve plant N nutrition, the engineering of biological nitrogen fixation (BNF) directly into cereal crops has been proposed as a future alternative to the overuse of chemical fertilizers (Bloch *et al*., 2020; Buren and Rubio, 2018; Curatti and Rubio, 2014; Oldroyd and Dixon, 2014).

BNF is the conversion of atmospheric dinitrogen gas into ammonia, a reaction catalyzed by the enzyme nitrogenase, which is found only in certain bacteria and archaea known as diazotrophs (Boyd *et al*., 2015; Dos Santos *et al*., 2012; Garcia *et al*., 2020). Recently, a unicellular eukaryotic alga containing a nitrogen-fixing organelle that evolved from a cyanobacterial endosymbiont has been reported (Coale *et al*., 2024). The most widespread form of nitrogenase is Mo-nitrogenase, which consists of two metalloproteins: MoFe protein (encoded by *nifD* and *nifK*) and Fe protein (encoded by *nifH*) (Bulen and LeComte, 1966; Spatzal *et al*., 2011). NifDK is a heterotetramer that harbors two types of complex iron-sulfur clusters in each half: an [8Fe−7S] P-cluster and an iron-molybdenum cofactor (FeMo-co) (Curatti *et al*., 2007; Shah and Brill, 1977). NifH is a homodimer that carries a single [4Fe–4S] cluster and acts as obligate electron donor to the MoFe protein (Seefeldt *et al*., 2020). Central to the maturation of nitrogenase is the assembly of its Fe−S cluster precursors carried out by NifU and NifS. NifU provides a scaffold for the assembly of [4Fe−4S] clusters, whereas NifS mobilizes sulfur (S) from cysteine for their synthesis on NifU (Smith *et al*., 2005). These clusters are then delivered to target nitrogenase proteins such as NifH and NifB (Dos Santos *et al*., 2004; Zhao *et al*., 2007). In *Azotobacter vinelandii*, a model diazotroph, deletion of either *nifU* or *nifS* abolishes nitrogenase activity, and cannot be rescued by the housekeeping ISC pathway for [Fe–S] cluster assembly (Dos Santos *et al*., 2007; Johnson *et al*., 2005).

The assembly of functional Mo-nitrogenase in a heterologous host must address two major challenges (Curatti and Rubio, 2014). The first one is the complexity of the nitrogenase biosynthetic pathway, which requires the co-expression of roughly 20 *nif* genes in precise stoichiometry to achieve an enzyme with optimal function (Buren *et al*., 2020; Buren *et al*., 2017; Poza-Carrion *et al*., 2014; Smanski *et al*., 2014; Temme *et al*., 2012; Yang *et al*., 2018). Second, nitrogenase [Fe–S] clusters and their biosynthetic intermediates are extremely sensitive to molecular oxygen (Eady *et al*., 1972; Shah and Brill, 1973).

The expression of nitrogenase components in yeast and plants has progressed considerably over the past decade. The earliest successes came from yeast, where *A. vinelandii* NifH expressed with NifM in mitochondria was active without any bacterial NifU or NifS, indicating that the endogenous ISC pathway was adequate for this protein (Lopez-Torrejon *et al*., 2016). However, producing a functional NifB in yeast mitochondria required co-expression with NifU, NifS and FdxN, probably because NifB consumes [4Fe–4S] clusters as substrates to generate NifB-co, a precursor to FeMo-co (Buren *et al*., 2019).

Experiments in tobacco have also proven the importance of NifU and NifS expression in chloroplasts. Endogenous SUF machineries can, to a limited extent, substitute for NifU and NifS in delivering [4Fe– 4S] clusters to NifH, as evidenced by the low but detectable NifH activity in transplastomic tobacco plants expressing only *nifH* and *nifM* under reduced atmospheric oxygen levels (Aznar-Moreno *et al*., 2021; Ivleva *et al*., 2016). When *nifU* and *nifS* from *A. vinelandii* were co-expressed with *nifH* and *nifM* and targeted to tobacco chloroplasts, a marked increase in Fe protein activity was observed (Eseverri *et al*., 2020). Transgenic tobacco and rice expressing NifH from *Hydrogenobacter thermophilus* in mitochondria yielded soluble proteins with low catalytic activity. However, its activity could be reconstituted *in vitro* by NifU and NifS, suggesting that the plant mitochondrial ISC pathway could not adequately deliver the NifH [4Fe-4S] cluster (Baysal *et al*., 2022; Jiang *et al*., 2021). The same pattern was observed for archaeal NifB from *Methanocaldococcus infernus* and *Methanothermobacter thermautotrophicus* expressed in transgenic rice mitochondria: soluble but cluster-depleted proteins that only became active after in vitro NifU reconstitution (He *et al*., 2022; Jiang *et al*., 2022). Taken together, these studies indicated that to achieve a functional nitrogenase in crops, the expression of functional NifU and NifS proteins was required. They also suggested that the previously used *A. vinelandii* NifU and NifS proteins were not optimal for this task.

Here, we describe the construction and screening of a library of NifU and NifS proteins from phylogenetically diverse organisms, their evaluation for chloroplast-targeting efficiency in rice protoplasts, and the stable integration of the three most promising NifUS pairs screened from *A. vinelandii* (Av), *Fischerella thermalis* (Ft), and *Marinobacterium lutimaris* (Ml) into the rice genome via *Agrobacterium tumefaciens*-mediated transformation. We further report the purification of NifU variants from agroinfiltrated *N. benthamiana* leaves, the *in vitro* reconstitution of [4Fe–4S] clusters on NifU, and the activation of apo-NifH by the transfer of NifU [4Fe–4S] clusters. Finally, we present a phenotypic characterization and differential proteome analysis in leaves of three-week-old transgenic rice plants expressing each NifUS pair, providing insights into rice developmental responses to the expression of NifU and NifS in chloroplasts.

## Materials and methods

### Bioinformatic search and sequence optimization of NifU and NifS variants

A list of 1003 organisms containing the minimum set of genes to infer diazotrophic activity (NifHDBKEN) from (Koirala and Brozel, 2021) was established as the starting point for library construction. The InterPro database was used to obtain a list of organisms that express NifU and NifS by searching for the corresponding entries (IPR016217: nitrogen fixation NifU; IPR017772: bacterial-type cysteine desulfurase NifS; and IPR017773: proteobacteria cysteine desulfurase NifS). Both lists were combined to detect organisms containing the NifHDKBEN and NifUS genes (358). Because Nif proteins from thermophilic organisms have been successfully expressed in eukaryotes (Buren *et al*., 2019; Jiang *et al*., 2022; Jiang *et al*., 2021; Meile *et al*., 2025) and nitrogenase is highly sensitive to oxygen, NifUS homologs from thermophilic and aerobic microbes were considered high-priority candidates. To identify them, the NCBI Taxon ID was used to obtain information on the growth temperature, habitats, and metabolism from diverse databases, including BacDive, JGI IMG/D, and GOLD, as described in (Meile *et al*., 2025).

The *nifU* and *nifS* sequences were codon-optimized for rice using DNA Chisel (Zulkower and Rosser, 2020). Optimization constraints were high GC (60%) and GC3 (75%) contents to increase expression (Sidorenko *et al*., 2017), avoidance of *Bpi*I, *Bsa*I, and *Bsm*BI restriction sites for cloning purposes, rare codons (minimum frequency=0.25), hairpins (stem size=15, window=200), and instability motifs, as previously reported (Eseverri *et al*., 2020). Optimized DNA sequences flanked by *Bpi*I sites for cloning were obtained from GenScript (USA).

### Generation of genetic constructs

Modular Cloning (MoClo) (Weber *et al*., 2011; Werner *et al*., 2012) was performed as described previously (Eseverri *et al*., 2020). To domesticate Level 0 parts, Phusion Hot Start II DNA Polymerase (Thermo Fisher) was used with the primers and templates listed in Supplementary Table S1. All generated constructs were verified by Sanger sequencing (Macrogen, Madrid, Spain) or by Oxford Nanopore whole-plasmid sequencing (Plasmidsaurus, Inc., Eugene, OR, USA). All synthetic genes, constructs, and the individual parts used for their assembly are listed in Supplementary Table S1.

### Rice protoplast transformation

Rice seeds (*Oryza sativa* L. cv. EYI 105) were sown in a 3:1 mixture of soil (Floragard, Oldenburg, Germany) and vermiculite in ⌀ 13 cm pots and grown for 9 days after germination in an environmentally controlled chamber (Conviron, Winnipeg, Canada) under 16/8 h light/dark conditions at 24 °C and 70% relative humidity. Protoplasts were isolated from rice seedlings as described by (Page *et al*., 2019) with slight modifications. Approximately 80 seedlings were harvested, and 10 cm of stem and sheath green tissue was retained and subsequently cut into 0.5–1 mm strip using a razor blade. The strips were immediately submerged in 0.6 M D-mannitol (Acros Organics) for 15 min in the dark at room temperature (RT) to initiate plasmolysis. After discarding the mannitol, the tissue was transferred to an enzyme solution (1.5% (w/v) cellulase RS (Duchefa), 0.75% (w/v) macerozyme R10 (Duchefa), 0.6 M D-mannitol (Acros Organics), 20 mM MES pH 5.7 (Duchefa), 10 mM KCl (Acros Organics), 10 mM CaCl2 (Merk), and 0.1% (w/v) BSA (GE Healthcare Life Sciences), and incubated in the dark for 4 hours at RT with 70 rpm shaking to allow digestion of cell wall material. An equal volume of W5 solution (154 mM NaCl, 125 mM CaCl2, 5 mM KCl, and 2 mM MES pH 5.7) was added to terminate the digestion.

Protoplasts were released by filtering through 40 µm nylon cell strainers (Corning) with 3 washes of the strips using W5 solution. The pellets were collected by centrifugation at 300 × *g* for 3 min at RT. The supernatant was decanted, and the pellets were washed by gentle resuspension in 10 mL of W5 solution. After the second centrifugation at 300 × *g* for 3 min at RT, the pellets were resuspended in 1–2 mL MMG solution (0.4 M D-mannitol, 15 mM MgCl_2_, and 4 mM MES pH 5.7). Protoplast transformation was performed by DNA-PEG-calcium transfection. For each transformation, 10 µL of plasmid (5–10 µg DNA) and 60 µL of protoplasts were combined with 70 µL of a freshly prepared PEG solution containing 40% (w/v) PEG 4000 (Merck, Darmstadt, Germany), 0.2 M D-mannitol, and 0.1 M CaCl_2_, and incubated in the dark at room temperature for 25 min. Negative control samples were prepared by replacing the plasmid with 10 µL of Milli-Q H_2_O. Next, 280 µL of W5 solution was slowly added to complete the transformation. Protoplasts were pelleted by centrifugation, resuspended in 375 µL of WI solution (0.5 M D-mannitol, 20 mM KCl, and 4 mM MES pH 5.7), transferred to 96-well microplates (125 µL per well), and incubated at room temperature for 16 h.

### Protein extraction and immunoblotting

Rice transformed protoplasts were recovered from 96-well plates by gentle pipetting, transferred to 1.5 mL Eppendorf tubes, and pelleted by centrifugation at 300 × *g* for 6 min at RT. The supernatants were discarded, and the pellets were shock-frozen in liquid nitrogen five times. To obtain total protein, pellets were resuspended in 2x Laemmli buffer, boiled for 10 min, cooled, and centrifuged at 14,000 × *g* for 3 min at 4 °C. To obtain the soluble protein fraction, pelleted protoplasts were resuspended in Protein Extraction Buffer (100 mM Tris-HCl pH 8, 200 mM NaCl, and 10% glycerol), incubated for 15 min on ice, centrifuged at 14,000 × *g* for 10 min at 4 °C, and the supernatant collected. To obtain the insoluble fraction, the previous pellet was washed several times with Protein Extraction Buffer to remove any remaining soluble protein, and resuspended in 2x Laemmli buffer. SDS-PAGE and immunoblot analyses were performed using standard methods. Immunoblots were visualized using an iBright FL1000 (Thermo Fisher Scientific) in chemiluminescence mode. Transgenic rice protein extracts were prepared from the aerial tissues of T0 plants to assess the expression and solubility of NifU and NifS variants by immunoblot analysis.

### Confocal microscopy

Samples were imaged using a Zeiss LSM880 confocal laser scanning microscope equipped with Plan-Apochromat 40X/1.2 water-immersion or Plan-Apochromat 63X/1.2 oil-immersion objectives. The excitation/emission wavelengths used with PMT detectors were as follows: GFP (488 nm/493–556 nm) and chlorophyll autofluorescence (633 nm/647–721 nm). Images were post-processed using ZEN 2.6 Blue (Zeiss) and ImageJ.

### Transgenic rice generation

Transgenic rice plants expressing selected NifU and NifS variants were obtained as described by (Hiei and Komari, 2008) with minor modifications. Embryogenic calli were induced from seeds of *Oryza sativa* L. cv. EY1 105 Japonica. After 4 weeks of culture, the calli were co-cultivated for 3–4 days with *Agrobacterium tumefaciens* carrying the constructs encoding the proteins of interest and then transferred to selective medium. Transformed calli were selected based on hygromycin resistance and GFP fluorescence included in the multigene constructs visualized in an iBright FL1000 in fluorescence mode. Positive calli were subsequently induced for shoot and root, and for plant regeneration. Once the regenerated plants (T0) reached an appropriate size and developmental stage, they were transferred to a greenhouse to continue growing and produce seeds for the next generation (T1). These seeds were used to obtain T2 plants for proteomic and phenotype characterization.

### Tobacco agroinfiltration

*N. benthamiana* plants were cultivated in 8 x 8 x 8 cm pots in a greenhouse in long-day conditions (16 h light/8 h dark) ensuring enough light with LED lamps at 26/22 °C day/night. Overnight cultures of *A. tumefaciens* GV3101 carrying the plasmid of interest were diluted to an OD_600_ of 0.8-0.9 in an infiltration solution containing 10 mM MES pH 5.5, 10 mM MgSO_4_, and 150 μM acetosyringone (Sigma-Aldrich) and incubated at 150 rpm at 24 °C for 3 h. Leaves of 3-4 weeks-old *N. benthamiana* plants were agroinfiltrated using a syringe without a needle and 3-4 days after infiltration, tissue was collected at the end of the dark period and used for protein extraction, confocal microscopy, and protein purification (protocols performed under ambient light).

### Purification of NifU from tobacco

Agroinfiltrated *N. benthamiana* leaves were frozen in liquid nitrogen and processed using an Oster Classic blender 4655 inside an anaerobic Coy Laboratory Glove box. Leaves were homogenized with lysis buffer (Tris–HCl 100 mM pH 8.6, NaCl 300 mM, glycerol 10%, PMSF 1 mM, leupeptin 1 µg/mL, DNase I 5 µg/mL in a 1:2 ratio). The extract was filtered through a cloth and centrifuged at 53,343 × *g* for 1.5 hour at 4° C. The supernatant was filtered through a Nalgene filter unit with a 0.2 µm pore size to obtain the cell-free extract (CFE). Twin-Strep (TS)-tagged NifU was purified by Strep-Tactin XT affinity chromatography under anaerobic conditions (< 0.1 ppm O_2_) using an AKTA Prime FPLC (GE Healthcare, Chicago, IL, USA) in an MBraun Glove box. The CFE was loaded into a 5-mL Strep-Tactin-XT high-capacity column (IBA Lifesciences) at a flow rate of 2.5 mL/min after equilibration with 20 column volumes of Buffer A (100 mM Tris–HCl pH 8, 300 mM NaCl, 10% glycerol), washed with 15 column volumes of Buffer A, and eluted in Buffer A supplemented with 50 mM biotin (IBA Lifesciences). Eluted protein was concentrated to 1.5 mL in an Amicon Ultra 30 K Centrifugal Filter unit (Merck Millipore, Burlington, MA, USA) with a 30 kDa cut-off pore size and desalted using a Sephadex G-25 PD-10 column (GE Healthcare, Chicago, IL, USA) to remove biotin. The elution from the PD-10 column was concentrated to 1-1.5 mL and stored in cryogenic tubes (Nalgene, Rochester, NY, USA) under liquid nitrogen.

### In vitro reconstitution of NifH activity with NifU

Apo-NifH was prepared *in vitro* according to (Rangaraj *et al*., 1997) with minor modifications. NifH purified from *A. vinelandii* (50 µM) was incubated with 40 mM α, α′-bipyridyl, 2.5 mM MgATP, and 2 mM sodium dithionite (DTH) in 22 mM Tris–HCl pH 7.4 for 30 min at 25 °C under anaerobic conditions. The resulting apo-NifH was desalted twice using PD-10 columns equilibrated with 100 mM Tris–HCl pH 8.0, 200 mM NaCl, 10% glycerol, and 2 mM DTH. Desalted apo-NifH was concentrated using an Amicon Ultra 30 K centrifugal filter and quantified using the BCA protein assay (Thermo Fisher Scientific, Waltham, MA, USA). The complete removal of NifH [4Fe–4S] cluster was confirmed by the loss of NifH acetylene reduction activity.

*In vitro* [4Fe–4S] cluster reconstitution of NifU variants was performed as described (Dos Santos *et al*., 2004; Lopez-Torrejon *et al*., 2016) with minor modifications. Each NifU variant isolated from tobacco (20 µM) was incubated in 22 mM Tris–HCl pH 7.4 containing 9 mM DTT for 30 min at 37 °C, followed by the addition of 0.4 mM (NH₄)₂Fe(SO₄)₂, 1 mM L-cysteine, and 225 nM pure *A. vinelandii* NifS. To avoid iron precipitation, (NH₄)₂Fe(SO₄)₂ and L-cysteine were added stepwise, in eight additions with 10–15 min incubations in between. Reconstitution reactions were incubated overnight at room temperature under anaerobic conditions. After reconstitution, DTT, free Fe, and L-cysteine were removed by buffer exchange using PD-10 columns pre-equilibrated with assay buffer.

The transfer of the [4Fe–4S] cluster from NifU to apo-NifH and the reconstitution of nitrogenase activity were simultaneously assessed using the acetylene reduction assay (ARA). Anaerobic reaction mixtures (400 µL final volume) contained purified *A. vinelandii* NifDK (0.22 µM), apo-NifH (8.73 µM), and NifU (87 µM) in an ATP-regenerating mixture consisting of 1.23 mM ATP, 18 mM phosphocreatine disodium salt, 2.2 mM MgCl₂, 3 mM DTH, and 46 µg/mL creatine phosphokinase in 22 mM MOPS buffer, pH 7.5. The molar ratios of NifDK:NifH and NifH:NifU were 1:40 and 1:10, respectively. Two positive control reactions were included: a holo-NifH purified from *A. vinelandii* (8.73 µM) replacing apo-NifH, and as-isolated [4Fe–4S] cluster-loaded NifU purified from *A. vinelandii* replacing the NifU variants. A negative control omitting NifU was included to confirm that apo-NifH alone did not support acetylene reduction. Reactions were transferred to 9-mL rubber-sealed vials under an argon atmosphere and initiated by injection of 1 mL acetylene (6% v/v final). After 20 min of incubation at 30 °C with shaking, reactions were stopped by addition of 100 µL of 8 M NaOH. Ethylene formation was quantified by gas chromatography using a Shimadzu GC-2014 fitted with a flame ionization detector (FID) and a Porapak N 80/100 column (Agilent Technologies, Santa Clara, CA, USA), with N₂ as carrier gas at 25 mL/min. Nitrogenase activity is expressed as nmol C₂H₄ · min⁻¹ · mg⁻¹ NifDK.

### Ultraviolet-visible spectroscopy (UV-vis)

Anaerobic protein samples for UV-vis were prepared inside an MBraun glovebox. Typically, 10-100 µL of NifU (as isolated or [Fe-S] cluster reconstituted) was added to 800 µL of Buffer A. Samples were transferred to sealed quartz cuvettes and scanned from 225 to 800 nm using a dual-beam Shimadzu UV-2600 spectrophotometer. Recorded absorbances were corrected by subtracting the sample absorbance at 800 nm, and spectra were normalized according to absorbance at 280 nm. The absorption spectra of air-oxidized proteins were recorded after bubbling the protein solution with air.

### Proteomic analysis

Three-week-old transgenic rice leaves expressing NifU and NifS were collected, and total protein was extracted using Protein Extraction Buffer 1:1 (v:w) and 2x Laemmli Buffer as described above for immunoblot analysis. Pools of T2 plants were used to generate four samples for each transgenic line (*Av*, *Ft*, and *Ml*) and four samples for wild-type rice plants.

Differential label-free quantitative (LFQ) proteomic analysis was conducted at the Proteomics Facility of Centro Nacional de Biotecnología (CNB, Madrid). Protein samples were digested under reducing and alkylating conditions using S-trap columns. Proteins were digested with 2 µg of trypsin, and the resulting peptides were purified and quantified using a Qubit fluorometer. Peptide yields were typically between 25 and 30 µg per sample. All peptide samples were adjusted to a final concentration of 0.2 µg · μL⁻¹, and 500 ng of peptides were injected for LC-MS/MS analysis. Peptide separation was performed by nano-liquid chromatography coupled with tandem mass spectrometry (nanoLC-ESI-MS/MS) on a reversed-phase C18 µPAC column using a 120 min chromatographic gradient and analyzed on an Orbitrap Exploris 240 mass spectrometer operating in Data Independent Acquisition (DIA) mode with variable isolation windows. Raw MS data were processed using Spectronaut v20.4 (Biognosys AG). Protein identification was performed using the *O. sativa* proteome database UP000059680_2026_03_17 downloaded from UniProtKB (17/03/2026), supplemented with a list of common laboratory contaminants (including albumin, keratins, and caseins). Spectronaut extracted peptide signals, assigned peptides to proteins using the razor + unique peptide approach, and performed data normalization. Protein abundance values were calculated from peptide intensities using Perseus (v1.6.15), and possible false positives were discarded using the following filters: proteins identified by at least two unique peptides, and proteins present in 75% of the replicates in at least one condition. Data were log2-transformed and missing values were imputed from a normal distribution (width = 0.3; down shift = 1.8). Differential protein accumulation between experimental conditions was assessed using an unpaired Student’s t-test, with p-values adjusted by permutation-based false discovery rate (FDR) correction (FDR=0.05; 250 randomizations), and differentially expressed proteins with a q-value < 0.05 were further analyzed.

### Phenotype characterization

Plant phenotyping was conducted using the automated Enclosed Ecosystem Phenotyping Platform (E2P2, LemnaTec) at the Centro de Biotecnología y Genómica de Plantas (CBGP, Madrid). Image acquisition was performed by capturing the lateral views of the plants under multiple wavelengths ranging from the blue spectrum to near infrared (NIR). Image analysis was performed using LemnaTec software. Plant segmentation was initially performed through color thresholding, followed by sequential image processing steps including noise reduction (denoising), erosion, and dilation to refine the segmentation. Statistical analyses were conducted using analysis of variance (ANOVA), followed by Tukey’s honestly significant difference (HSD) test as a post hoc analysis, considering a significance threshold of p < 0.05. When the assumption of homoscedasticity was not met, a non-parametric Kruskal–Wallis test was applied, followed by Dunn’s test for multiple comparisons, using a significance threshold of p < 0.05.

## Results

### Selection of NifU and NifS variants

To identify NifU and NifS variants with enhanced performance in plant cells, homologs were selected from six phylogenetically diverse diazotrophic organisms representing distinct ecological niches and physiological properties (Supplementary Fig. S1). The main selection criteria for the final candidates of the library were high growth or isolation temperature, aerobic growth, extreme habitats, and taxonomic diversity. These criteria aimed to increase the probability of identifying proteins with enhanced stability and robustness within the plant cellular environment, as well as an enhanced capacity to support [Fe–S] cluster assembly and transfer to downstream components of the nitrogenase pathway, including the NifH protein tested here. The super index in NifU and NifS proteins described in this work denotes the organism of origin. The NifH and NifDK proteins used in biochemical analysis are from *A. vinelandii*.

### Chloroplast targeting and solubility of NifU and NifS variants in rice protoplasts

NifU and NifS variants were fused to chloroplast transit peptides (CTPs) for chloroplast targeting and expressed in rice protoplasts (Supplementary Fig. S2A). Initially, CTPs previously validated for the *A. vinelandii* NifU*^Av^* and NifS*^Av^* (Eseverri *et al*., 2020) were employed. When these CTPs were applied to the newly selected NifU and NifS, several variants run as multiple bands in SDS-PAGE and immunoblot analysis (Supplementary Fig. S2B). This pattern typically indicates incomplete processing of the CTP resulting in suboptimal targeting or maturation of the affected protein. Thus, alternative CTPs were used to identify those that produced the expected processed protein size for each NifU and NifS homolog in immunoblot analysis (Supplementary Fig. S2C-D). Correct processing was substantiated by comparing the apparent molecular weights of chloroplast-targeted NifU and NifS proteins with those of their cytosolic counterparts lacking transit peptides (Fig. 1). Fractionation analysis of protein extracts from rice protoplasts were performed in parallel and revealed differences in solubility among homologs, with some variants predominantly present in the soluble fraction, while others were also present in the insoluble fraction (Supplementary Fig. S3). Processed proteins present in the soluble fraction and matching the expected size following transit peptide cleavage were selected. For these, the subcellular localization to the chloroplast stroma was further assessed by confocal microscopy (Fig. 2). The *A. vinelandii*, *F. thermalis*, and *M. lutimaris* NifU and NifS met the requirements of expression, solubility, CTP processing, and chloroplast localization, and were suitable for functional testing in planta.

**Fig. 1.**
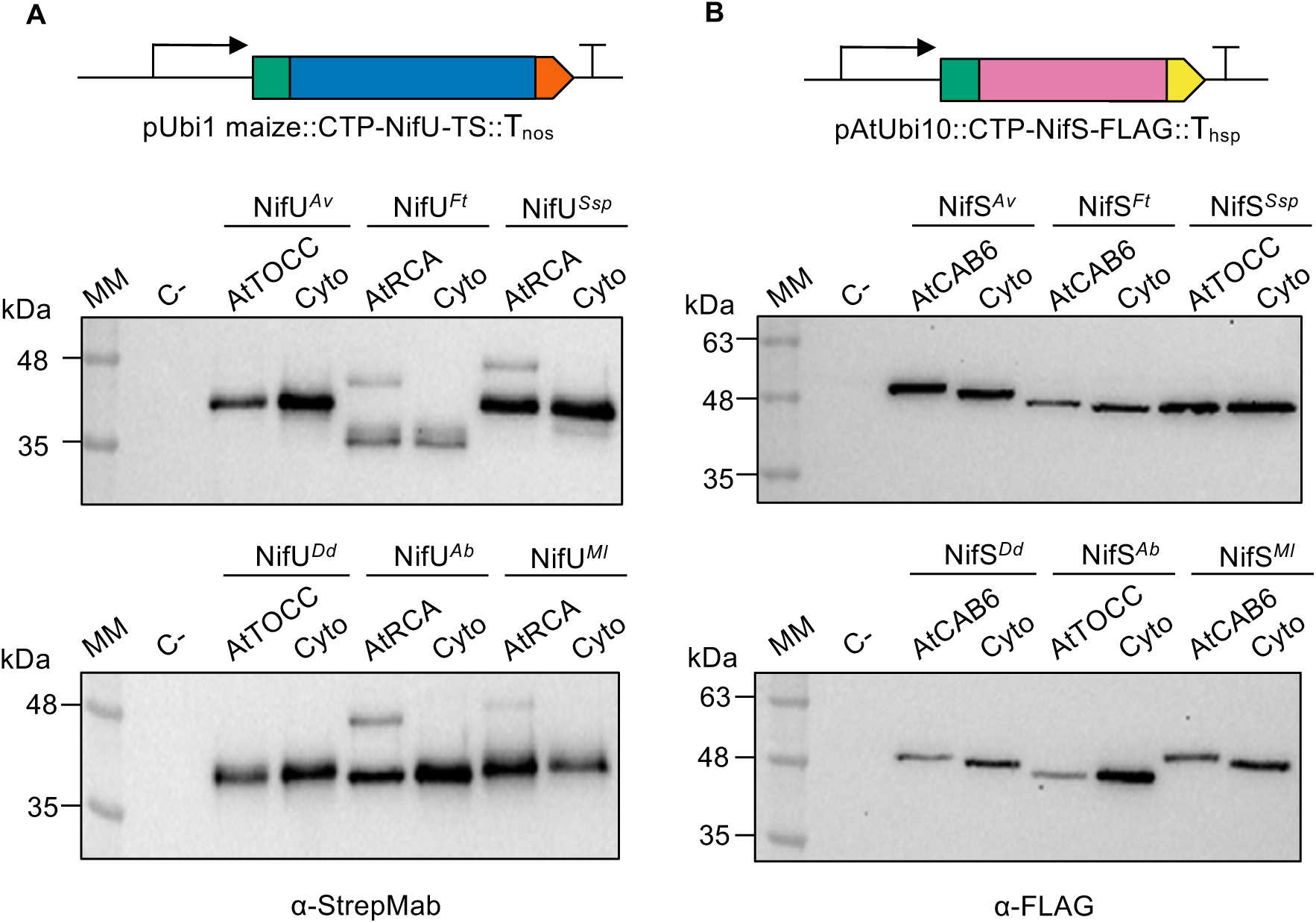
Rice protoplast CTP selection for NifU (A) and NifS (B) variants. SDS-PAGE and immunoblot analysis performed with proteins extracts from protoplasts transformed with the *nifU* and *nifS* genes either fused to a CTP (AtTOCC, AtRCA, or AtCAB6) or lacking the CTP for cytosolic localization. C-refers to protein extracts from non-transformed rice protoplasts. Species abbreviations: *Azotobacter vinelandii* (*Av*), *Fischerella thermalis* (*Ft*), *Synechococcus sp.* JA-2-3B’a (*Ssp*), *Dechloromonas denitrificans* (*Dd*), *Azospirillum baldaniourum* (*Ab*), and *Marinobacterium lutimaris* (*Ml*). Graphic representations of the genetic constructs are shown at the top of each panel: CTP (green), Twin-Strep tag (TS), orange, NifU (blue), NifS (pink), and FLAG tag (yellow).

**Fig. 2.**
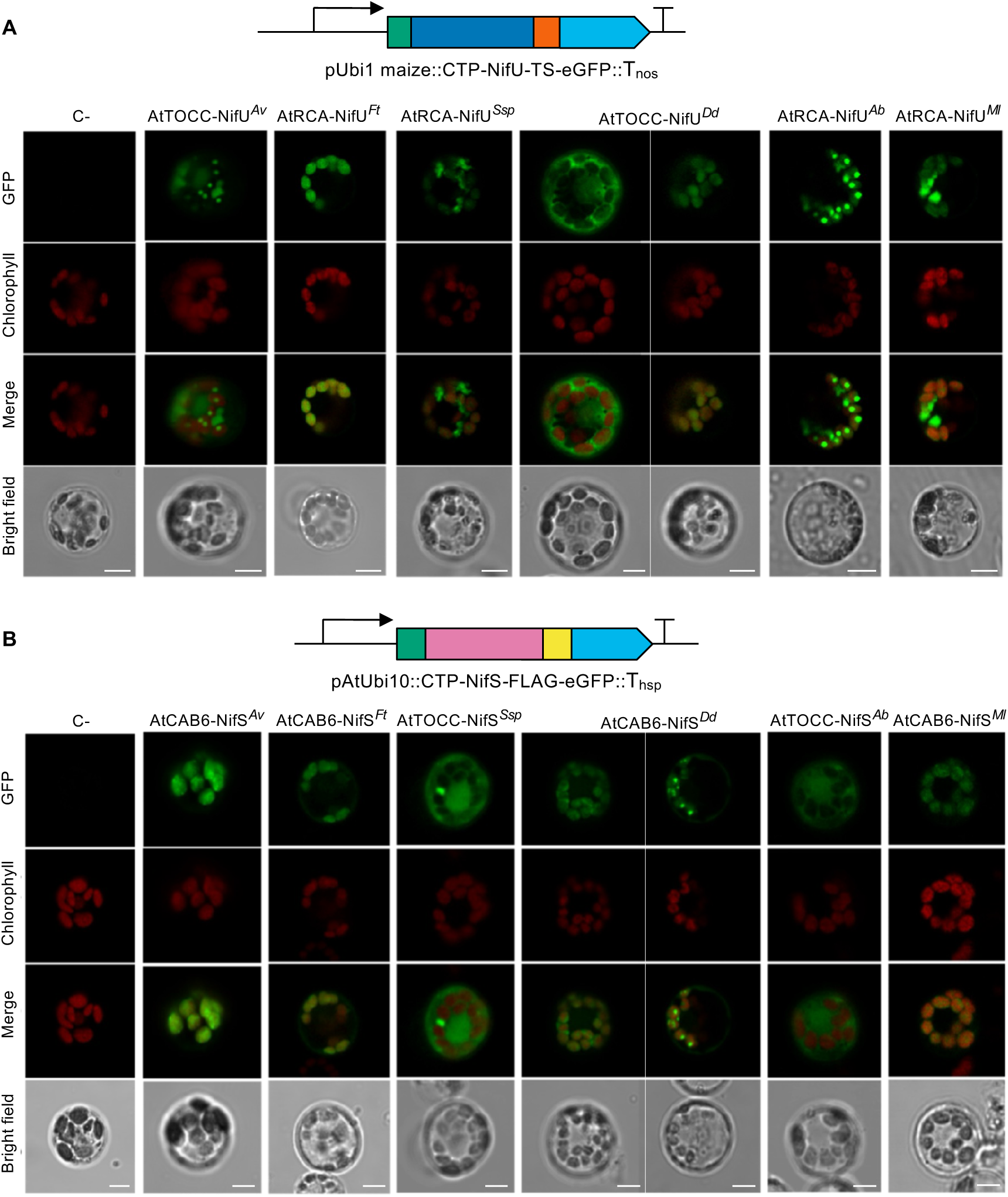
Confocal microscopy images of the subcellular localization of NifU (A) and NifS (B) variants. NifU and NifS variants were expressed in rice protoplasts fused to a CTP and the green fluorescent protein (GFP). Species abbreviations as in Fig.1. Co-localization with chlorophyll autofluorescence (third row, merge) indicates correct import of NifU and NifS variants from *F. thermalis* and *M. lutimaris* into chloroplasts. Scale bars, 5 µm. C-: non-transformed rice protoplasts. Graphic representations of the genetic constructs are shown at the top of each panel: CTP (green), Twin-Strep tag (TS), orange, NifU (blue), NifS (pink), FLAG tag (yellow), and eGFP (cyan).

### Function of NifU and NifS variants purified from N. benthamiana

*Agrobacterium*-mediated transient expression in *N. benthamiana* leaves was employed for rapid and scalable protein production, as rice protoplasts do not support protein accumulation at the scale required for purification. Immunoblot and confocal microscopy analyses confirmed that the selected NifU and NifS variants were expressed, targeted to the *N. benthamiana* chloroplasts and correctly processed (Supplementary Fig. S4), suggesting that the rice protoplast validation protocol was robust and transferable to other plant species.

The three TS-tagged NifU variants were purified by Strep-Tactin affinity chromatography (Supplementary Fig. S5A). As-isolated, NifU variants contained low levels of [Fe–S] clusters according to the intensity of characteristic 320 nm and 400-420 nm absorption bands in the UV-vis spectra (Supplementary Fig. S5B). However, NifU variants readily acquired the expected spectral signatures upon *in vitro* [Fe–S] cluster reconstitution, indicating retention of the structural integrity required for [Fe–S] cluster assembly. Exposure of reconstituted NifU proteins to air changed UV-vis spectra as expected for O_2_ sensitive [Fe–S] clusters (Supplementary Fig. S5C). Together, these results suggest limitation or instability of [Fe–S] cluster loading in planta (or losses during purification) rather than intrinsic defects of produced NifU proteins.

The [Fe–S] cluster loaded NifU variants were tested for their ability to reconstitute activity of [4Fe–4S] cluster-deficient apo-NifH*^Av^* in its *in vitro* reaction with NifDK*^Av^*. Apo-NifH*^Av^* lacked measurable activity without reconstitution, and non-reconstituted NifU proteins were unable to activate it (n = 3). Reconstituted NifH*^Av^* activities (n = 3) were 1497 ± 12.48, 855 ± 60.95, and 413 ± 39.20 nmol ethylene formed·min⁻¹·mg⁻¹ NifDK when activated by NifU*^Ft^*, NifU*^Av^*, and NifU*^Ml^*, respectively. In comparison, the [Fe–S] cluster loaded NifU*^Av^* purified from *A. vinelandii* cultures produced 1108 ± 26.30 units of activity (n = 3). Thus, NifU proteins produced in planta were competent for [Fe–S] cluster assembly and apo-NifH*^Av^*activation *in vitro*.

### Phenotypic effects of stable NifU and NifS expression in rice plants

Transgenic rice plants constitutively expressing NifU and NifS either from *F. thermalis*, *M. lutimaris*, or *A. vinelandii* were generated using multigenic constructs (abbreviated as *Av*, *Ft* and *Ml* plants for simplicity) (Fig. 3A). Immunoblot analysis of protein extracts obtained from primary transformants (T0 generation) confirmed that NifU (Fig. 3B) and NifS (Fig. 3C) variants were soluble and exhibited apparent molecular weights consistent with correct chloroplast targeting and CTP processing, as previously validated in rice protoplasts.

**Fig. 3.**
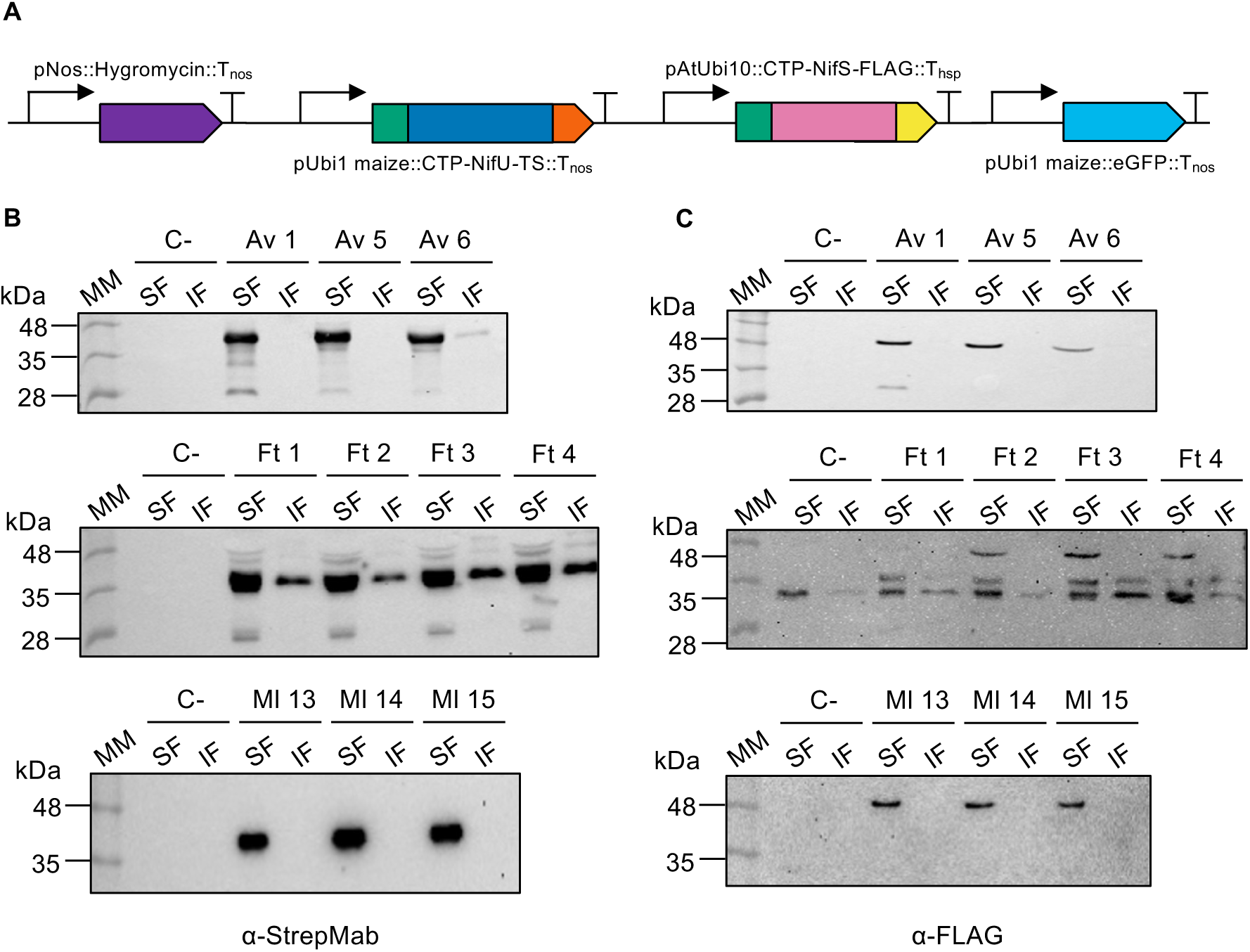
Solubility screening of NifU and NifS variants with selected CTP in transgenic regenerated rice plants. (A) Multigenic constructs were generated for each CTP-NifU and CTP-NifS pair (*A. vinelandii, F. thermalis* and *M. lutimaris*), including a hygromycin resistance cassette and a transcriptional unit for cytosolic GFP expression to select for positive transformants. SDS-PAGE and immunoblot analysis performed with soluble (SF) and insoluble (IF) fractions from leaf samples of T0 regenerated transgenic plant lines (Av, Ft, and Ml) developed with antibodies against TS-NifU (B), or FLAG-NifS (C). C-: soluble and insoluble fractions from wild-type rice leaves.

Homozygous T₂ lines expressing the different NifU and NifS variants exhibited consistent differences in plant growth and physiological status, with marked reduction in size and vigor being most apparent for *Av* plants followed by the *Ft* plants. In contrast, *Mt* plants were indistinguishable from the wild type (Fig. 4A). Quantitative analysis of growth parameters, such as leaf area (Fig. 4B) and spectral indexes associated with photosynthetic performance and pigment composition (NDVI, ND705, ARI1, and SIPI) (Fig. 4C and Supplementary Fig. S6) consistently suggested that growth defects were due to altered physiological status rather than solely developmental effects.

**Fig. 4.**
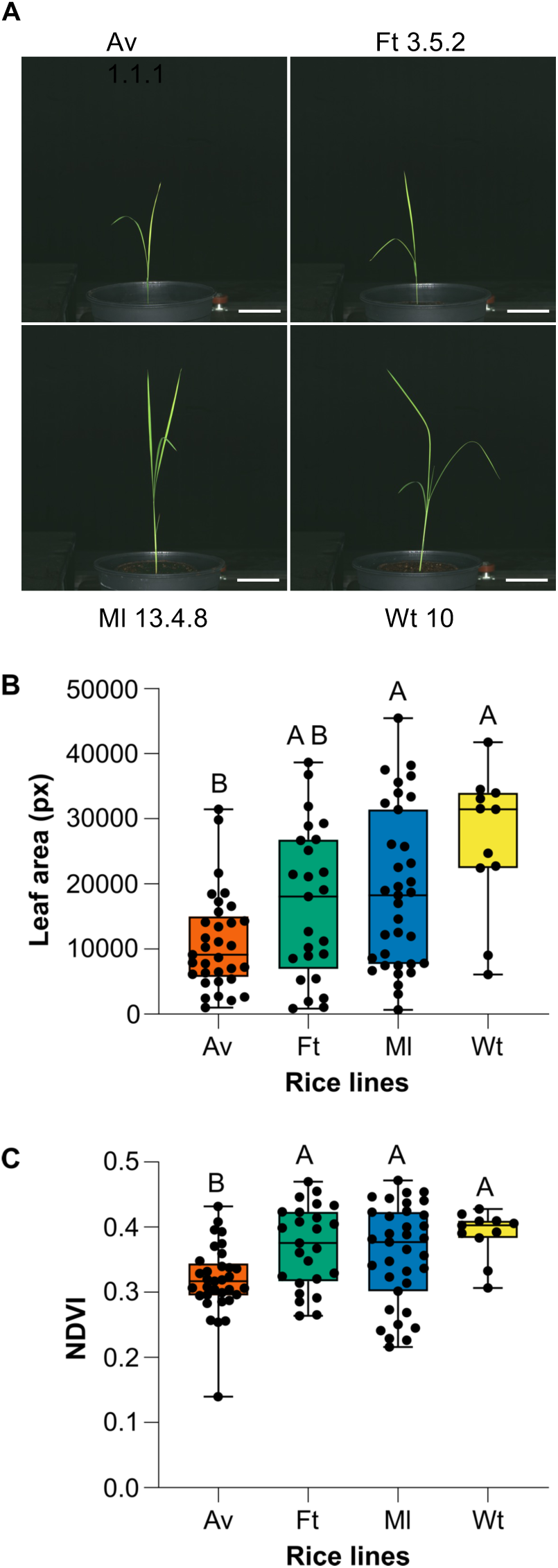
Phenotype of rice transgenic lines expressing NifU and NifS variants. (A) Three weeks-old wild-type and transgenic rice lines expressing *A. vinelandii* (*Av*), *F. thermalis* (*Ft*), or *M. lutimaris* (*Ml*) NifUS. Boxplots showing leaf area measurements (B), and Normalized Difference Vegetation Index (NDVI) (C) of wild-type and *Ml*, *Ft*, and *Av* plants. Each point represents an individual biological replicate. Boxes show interquartile range, the center line indicates the median, and whiskers represent the minimum and maximum values. Different letters indicate significant differences among phenotypes according to Kruskal–Wallis analysis followed by Dunn’s post hoc test (*p* < 0.05).

### NifU and NifS expression triggers variant-specific proteomic shifts

Comparative proteomics of rice seedlings was performed to investigate the molecular basis underlying phenotypic differences. We hypothesized that integration into endogenous plant pathways and disruption of host metabolism could be variant-dependent and have a proteome effect. Global analysis of differentially accumulated proteins revealed that expression of NifU and NifS variants predominantly triggered an increase in protein abundance. Both, the total number of overexpressed proteins and the maximum fold changes followed the order *Av* > *Ft* > *Mt* plants (Fig. 5).

**Fig. 5.**
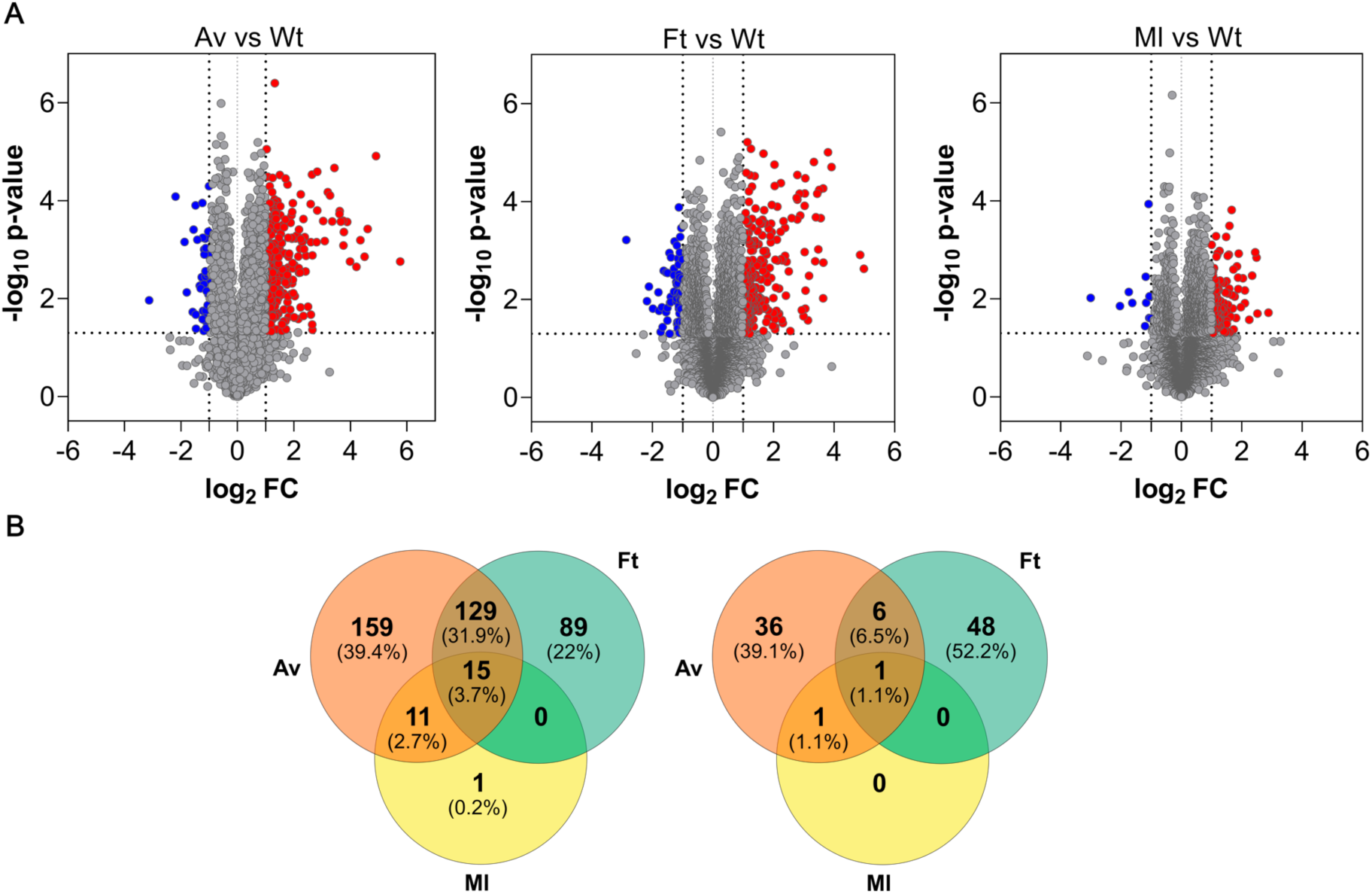
NifUS-variant dependent proteome changes in transgenic rice. (A) Volcano plots generated with Perseus based on a dataset of 5,700 proteins that passed the quality test (see Materials and Methods). Each plot compares the proteomes of *Av*, *Ft*, and *Ml* lines against the wild-type (*wt*) reference to evaluate up- and down-regulated proteins and their statistical significance. Each red dot on the right and blue dot on the left represent an up- or down-regulated individual protein in comparison with the *wt* and the stated threshold levels (dashed line). (B) Venn diagrams illustrating the quantitative overlap of proteins that were significant up- (left) or down-regulated (right) among the rice lines.

Pathway enrichment analysis of overexpressed proteins across all variants revealed that only 3.7% followed a common trend, namely the enhanced accumulation of serine protease inhibitors (B7E9D7, A5HEI2, Q0JR27) (Supplementary Table S2). In contrast, Nucleoredoxin 1 (NRX1; Q7Y0E8, Os03g0405500) was not retrieved by the G:Profiler algorithm under any enriched GO term but was consistently identified as strongly upregulated through direct inspection of individual fold changes (Supplementary Table S3). NRX1 showed an 11-fold increase in *Av* and *Ft* plants and a 4-fold increase in *Ml* plants. This gradient suggests a progressive oxidative stress response modulated by the expression of different NifUS variants consistent with NRX1 role as a central regulator of antioxidant defense and redox homeostasis, as it has been shown that mutants lacking NRX1 are hypersensitive to ROS (Kneeshaw *et al*., 2017).

The most pronounced change in the dataset was observed in *Av* plants, where 159 proteins (119 GO classified) were enriched in “hydrolase activity,” with 40 belonging to this category and 12 acting on O-glycosyl compounds. For example, BGLU30 (Q0J0N4) was 5-fold overexpressed in *Av* plants, consistent with its role in sulfur-deficiency induced glucosinolate catabolism (Zhang *et al*., 2020) (Supplementary Table S4).

Interestingly, 129 proteins (110 classified into GO terms) were overexpressed in *Av* and *Ft* but not in *Ml* plants. Of these, 82 mapped to “catalytic activity,” including 27 with “oxidoreductase activity” (Supplementary Table S5). Defense-related proteins accumulated strongly in *Av* and *Ft* plants. JAC1 (Q306J3) was overexpressed 56-fold in *Av* and 31-fold in *Ft*; lipoxygenase Q2QNN5 increased 19 and 29-fold; multiple chitinases (O22080, Q10S66, Q7XCK6, Q8RU26) rose 7 to 25-fold depending on the genotype. BBI-like protease inhibitors also accumulated significantly in *Av* and *Ft*, but not in *Ml*. Antioxidant enzymes such as glutathione transferases (Q8L4V6, Q93WY5, Q0JLY2), peroxidases (Q7F1U0, Q0DCP0, Q5JMS4), and ferritin were strongly induced, with fold changes ranging from 4-to over 13-fold. This response, which is consistent with activation of antioxidant defenses under conditions of metal ion imbalance (Fourcroy *et al*., 2004), was absent in *Ml* plants.

A very small fraction of proteins (0.3%) was detected exclusively in *Av* or *Ft* plants (Supplementary Fig. S7 and Table S6), suggesting that their abundance in the wild-type and *Ml* lines was below the detection threshold. Some of these proteins overlapped with GO terms that were significantly enriched in *Av* and/or *Ft* plants, such as peroxidases (Supplementary Tables S4 and S5).

Very few proteins were commonly downregulated across all variants, while 36 proteins (22–25 GO classified) were specifically reduced in *Av* plants. Of these, 13 mapped to “chloroplast,” six to “photosynthesis,” and four to [Fe–S] cluster binding (Supplementary Table S7). Closer inspection of chloroplast Fe–S proteins revealed variant-dependent impacts: some increased, others decreased, with *Ml* plants showing the least perturbation (Supplementary Fig. S8). This suggests interference with endogenous [Fe–S] cluster assembly and allocation, destabilizing apo-proteins when cluster insertion was deficient. In contrast, the accumulation level of proteins involved in chloroplast [Fe–S] cluster biosynthesis remained mostly unchanged, except for a subtle modification of SUFB (Q941V2), SUFC (Q10LW5), and SUFD (Q5ZDZ8), and a ∼30% reduction in chloroplast GRXS12 (Q2QX01), a key [Fe–S] cluster transfer protein (Supplementary Fig. S9). Overall, changes in the relative levels of proteins of the mitochondrial biosynthetic pathway for [Fe-S] clusters was modest. However, expression of the Ft-NifUS variant produced a statistically significant increase of 4.4-, 1,3-, and 1.4-fold for HSCA3 (B9G4B3). HSCA1 (Q0DX50), and ISA3 (A3BSI8), respectively.

## Discussion

Introducing a functional nitrogenase pathway into plant cells remains a major challenge in plant biotechnology. A key step in this process is the successful reconstitution of the [Fe–S] cluster assembly pathway required to support nitrogenase function. This is exemplified by the nitrogenase component NifH and the nitrogenase cofactor biosynthetic protein NifB, which have been expressed in plant mitochondria and chloroplasts but have limited function due to insufficient [Fe–S] cluster content (Baysal *et al*., 2022; He *et al*., 2022; Jiang *et al*., 2022; Jiang *et al*., 2021). Earlier work showed that *A. vinelandii* NifU and NifS can be targeted to chloroplasts with suboptimal performance (Aznar-Moreno *et al*., 2021; Eseverri *et al*., 2020). Here, we extend these findings by comparing homologs from diverse diazotrophs and demonstrating that NifU/NifS variant selection critically influences compatibility with plant metabolism.

NifU proteins produced in plant cells are largely devoid of [Fe–S] clusters in the as-isolated state yet remain competent for *in vitro* reconstitution. This suggests that the proteins are correctly folded but inefficiently loaded with clusters in planta, pointing to [Fe–S] cluster availability as a major bottleneck. This limitation is consistent with the highly regulated nature of chloroplast [Fe–S] assembly, which is primarily devoted to endogenous client proteins, particularly those involved in photosynthetic electron transport (Balk and Pilon, 2011; Couturier *et al*., 2013). Consequently, heterologous proteins may have restricted access to this machinery, especially under high expression conditions.

A central finding of this work is that NifU and NifS proteins from different diazotrophic origins display markedly distinct behaviors when expressed in plant chloroplasts. Although all variants are correctly targeted, processed, and accumulated, their impact on [Fe–S] cluster metabolism, proteome profile, and plant physiology varies substantially. Despite efficient expression, *A. vinelandii* variants strongly impair plant growth and photosynthetic performance and trigger extensive proteome reprogramming, including downregulation of photosynthetic processes and induction of stress-related responses. *F. thermalis* variants produce intermediate effects, whereas *M. lutimaris* variants produce minimal perturbations of growth and physiology appearing compatible with the plant cellular environment. These observations highlight the importance of host compatibility in engineering nitrogenase pathways.

Phenotypic and proteomic data further support the idea that heterologous NifU and NifS expression interferes with endogenous chloroplast processes, particularly those associated with Fe–S metabolism. The NVDI index, which correlates with photosynthetic performance and chlorophyl content (Gamon *et al*., 1995), was severely affected in *Av* lines. The pronounced reduction in photosynthetic performance and selective downregulation of photosynthesis-related proteins observed in *Av* lines are indicative of impaired chloroplast function. Given that Photosystem I is highly enriched in [Fe–S] clusters essential for electron transfer, disruption of its assembly or maintenance provides a plausible explanation for the observed phenotypes (Schottler and Toth, 2014). Therefore, we propose that overexpression of NifU and NifS leads to competition with endogenous pathways for [Fe–S] cluster assembly or delivery, limiting cofactor availability for native proteins and ultimately compromising photosynthetic function.

This perturbation is accompanied by coordinated stress responses. The induction of JAC1, lipoxygenase, chitinases, and BBI-like endopeptidase inhibitors indicates activation of defense pathways, while increased abundance of glutathione transferases, peroxidases, ferritin, and NRX1 reflects a strong antioxidant response. These systems act to detoxify reactive oxygen species (ROS), regulate iron homeostasis, and maintain chloroplast redox balance. Notably, ferritin induction suggests altered iron metabolism, a process increasingly recognized as intertwined with plant immunity: iron sequestration and deficiency responses can activate defense signaling, as shown by the immune-stimulating activity of bacterial siderophores (Herlihy *et al*., 2020). Thus, activation of oxidative stress and immune responses, both related processes, are consistent with recombinant NifUS-dependent perturbation of redox balance and or iron metabolism in the transgenic plants. The gradient of NRX1 induction (*Av* > *Ft* > *Ml*) reflects the severity of stress imposed by each variant.

Simultaneously, modest but consistent alterations in the abundance of endogenous Fe–S proteins suggest a broader disruption of cofactor homeostasis. Most proteins decreased slightly, consistent with the instability of apo-forms when cluster supply was limited, whereas a few showed minor increases. This pattern resembles previous observations under Fe limitation (Crooks *et al*., 2010; Reyda *et al*., 2008) and supports the notion that heterologous NifUS expression perturbs Fe–S protein maturation. The absence of cognate apo-Nif proteins may further exacerbate these effects by diverting [Fe–S] precursors toward non-native targets.

HSCAs are molecular chaperones that specifically recognize a conserved sequence in IscU (bacteria) and ISU (mitochondria) (Vickery and Cupp-Vickery, 2007). In plants, both HSCA1 and HSCA3 are stress-responsive, being induced in rice under conditions such as oxidative stress and iron depletion (Liang *et al*., 2014). Thus, plant sensing of oxidative stress and iron imbalance due to recombinant NifUS in the chloroplast feeds back on mitochondrial [Fe–S] cluster biosynthesis. This interference exacerbates a self-sustaining cycle of metabolic disturbance consistent with a broader model in which disruption of [Fe–S] cluster assembly impairs electron transport and redox homeostasis, leading to electron leakage and ROS generation. These ROS, in turn, activate antioxidant and defense pathways that only partially offset the underlying imbalance. Subtle perturbations in components such as GRXS12 may amplify this instability, since reduced GRXS12 function has been linked to redox disequilibrium, defective Fe–S protein maturation, and chloroplast dysfunction in *Arabidopsis thaliana* (Liu *et al*., 2026). Overall, the intensity of this self-reinforcing process followed the gradient *Av* > *Ft* > *Ml*, underscoring variant-specific perturbations in redox and iron metabolism.

Interestingly, this study revealed some uncoupling between biochemical performance and physiological compatibility. Although *F. thermalis* NifU shows the highest capacity to activate *A. vinelandii* NifH *in vitro*, it still imposes a measurable physiological burden, whereas *M. lutimaris* NifU displays lower activity but minimal impact on plant growth. This trade-off highlights the importance of evaluating NifU variants within their functional context. Moreover, because the activation assays rely on *A. vinelandii* NifH, they may not reflect optimal pairing with alternative NifH proteins, such as the *Hydrogenobacter thermophilus* NifH protein successfully expressed in rice (Baysal *et al*., 2022). In addition, NifU participates in multiple steps of nitrogenase assembly, including FeMo-co biosynthesis via NifB, so its selection cannot be based solely on NifH activation capacity.

Overall, these results emphasize that the successful implementation of nitrogenase in plants will require more than just the transfer of individual genes. Interactions between heterologous components and endogenous metabolic networks, particularly those governing [Fe–S] cluster assembly, represent a major constraint. Rather than a single optimal solution, a network-conscious optimization strategy should be considered, in which NifU variants are selected according to their compatibility with specific NifH, NifDK, NifEN, and NifB partners, and host physiology. In this context, the strong compatibility of *M. lutimaris* NifU, despite its lower *in vitro* activity, and the balanced activity versus compatibility of *F. thermalis* NifU make them attractive candidates for further optimization. Therefore, future efforts should focus on evaluating combinations of Nif components while enhancing host [Fe–S] cluster assembly capacity, for example through modulation of the SUF pathway or the introduction of dedicated metal delivery and storage systems.

## Supplementary data

Fig. S1: Library of NifUS homologs.

Fig. S2. Screening of CTP to target NifU and NifS variants to rice chloroplasts.

Fig. S3. Solubility screening of NifU and NifS variants with selected CTP in rice protoplasts.

Fig. S4. Transient expression of selected NifU and NifS variants in *N. benthamiana* leaves.

Fig. S5. Biochemical characterizations of NifU variants purified from *N. benthamiana* leaves.

Fig. S6. Additional phenotype parameters of wild type (Wt) and transgenic *Ml*, *Ft*, and *Av* lines.

Fig. S7. Venn diagram showing the overlap of proteins identified in wild-type, *Av*, *Ft*, and *Ml* rice plants.

Fig. S8. Relative expression of [Fe-S] cluster proteins and ferritin in *Av*, *Ft* and *Ml* rice lines.

Fig. S9. Relative expression of [Fe-S] cluster biosynthesis proteins in *Av*, *Ft* and *Ml* rice lines.

Table S1. Genes, primers, and plasmids used in this study.

Table S2. G:Profiler pathway enrichment analysis of overexpressed proteins in *Av*, *Ft* and *Ml* rice lines.

Table S3. Top 20 most upregulated or downregulated proteins in transgenic lines.

Table S4. G:Profiler pathway enrichment analysis of downregulated proteins in *Av* rice lines (but not in *Ft* or *Ml* lines).

Table S5. G:Profiler pathway enrichment analysis of overexpressed proteins in *Av* and *Ft* rice lines (but not in *Ml* lines).

Table S6. Proteins detected only in specific *Av*, *Ft* or *Ml* rice lines.

Table S7. G:Profiler pathway enrichment analysis of downregulated proteins in *Av* rice lines (but not in *Ft* or *Ml* lines).

## Acknowledgements

We thank Carlos Echavarri for providing NifH and NifDK proteins.

## Author contributions

LC and LMR acquired funding; ME, EC, and LMR designed research; ME, CV, and NM-S, performed the experimental research; ME, AOS, and LC, worked with proteomic data; FB performed the phenotypic analysis; EC, LC, and LMR supervised work; All authors analyzed data and contributed to manuscript preparation.

## Data availability

All data supporting the findings of this study are available within the paper and its Supplementary Information. A preprint version of the manuscript is available at bioRxiv under a CC BY 4.0 license. The mass spectrometry proteomics data have been deposited to the ProteomeXchange Consortium via the PRIDE (Perez-Riverol *et al*., 2025) partner repository with the dataset identifier PXD080770.

## Abbreviations

CTP: chloroplast targeting peptide
DTH: dithionite
Av: *Azotobacter vinelandii*
Ft: *Fischerella thermalis*
Ml: *Marinobacterium lutimaris*

