## Supplemental Figures and Tables for "Natural variation in NifU and NifS enhances chloroplasts compatibility for nitrogenase engineering"

### Supplementary data

Fig. S1: Library of NifUS homologs.

Fig. S2. Screening of CTP to target NifU and NifS variants to rice chloroplasts.

Fig. S3. Solubility screening of NifU and NifS variants with selected CTP in rice protoplasts.

Fig. S4. Transient expression of selected NifU and NifS variants in *N. benthamiana* leaves.

Fig. S5. Biochemical characterizations of NifU variants purified from *N. benthamiana* leaves.

Fig. S6. Additional phenotype parameters of wild type (Wt) and transgenic *Ml*, *Ft*, and *Av* lines.

Fig. S7. Venn diagram showing the overlap of proteins identified in wild-type, *Av*, *Ft*, and *Ml* rice plants.

Fig. S8. Relative expression of [Fe-S] cluster proteins and ferritin in *Av*, *Ft* and *Ml* rice lines.

Fig. S9. Relative expression of [Fe-S] cluster biosynthesis proteins in *Av*, *Ft* and *Ml* rice lines.

Table S1. Genes, primers, and plasmids used in this study (Excel spreadsheets).

Table S2. G:Profiler pathway enrichment analysis of overexpressed proteins in *Av*, *Ft* and *Ml* rice lines.

Table S3. Top 20 most upregulated or downregulated proteins in transgenic lines.

Table S4. G:Profiler pathway enrichment analysis of downregulated proteins in *Av* rice lines (but not in *Ft* or *Ml* lines).

Table S5. G:Profiler pathway enrichment analysis of overexpressed proteins in *Av* and *Ft* rice lines (but not in *Ml* lines).

Table S6. Proteins detected only in specific *Av*, *Ft* or *Ml* rice lines.

Table S7. G:Profiler pathway enrichment analysis of downregulated proteins in *Av* rice lines (but not in *Ft* or *Ml* lines).

**A**

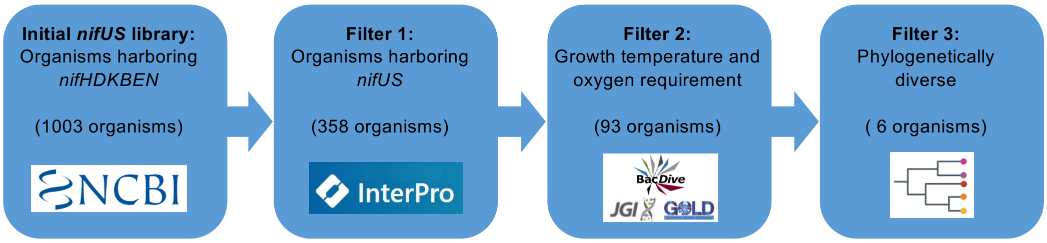


**B**

| Organism name (NCBI) | Phylum | Class | Order | Isolation | Oxygen Requirement | Optimal Growth Temperature (ºC) |
|---|---|---|---|---|---|---|
| *Azotobacter vinelandii* (strain DJ / ATCC BAA-1303) | Proteobacteria | Gammaproteobacteria | Pseudomonadales | Soil free-living | Aerobe | 28 |
| *Fischerella thermalis* CCMEE 5201 | Cyanobacteria | Unclassified | Nostocales | Thermal spring | Aerobe | 45-50 |
| *Synechococcus sp.* JA-2-3B'a(2-13) | Cyanobacteria | Unclassified | Synechococcales | Thermal spring | Facultative | 50-55 |
| *Dechloromonas denitrificans* ED1, ATCC BAA-841 | Proteobacteria | Betaproteobacteria | Rhodocyclales | Earthworm gut *(Aporrectodea caliginosa)* | Facultative anaerobe | 30 |
| *Azospirillum baldaniorum* (sp 245) | Proteobacteria | Alphaproteobacteria | Rhodospirillales | Rhizosphere | Aerobe | 30 |
| *Marinobacterium lutimaris* AN9, DSM 22012 | Proteobacteria | Gammaproteobacteria | Oceanospirillales | Tidal flat sediment | Aerobe | 25-30 |

**Fig. S1.** Library of NifUS homologs. (A) Schematic representation of filtering criteria showing the number of candidates passing each filter and the final selection. (B) List of organisms whose *nifUS* genes were included in the screening. The list contains relevant information about phylogeny, habitat, oxygen requirement, and optimal growth temperature.

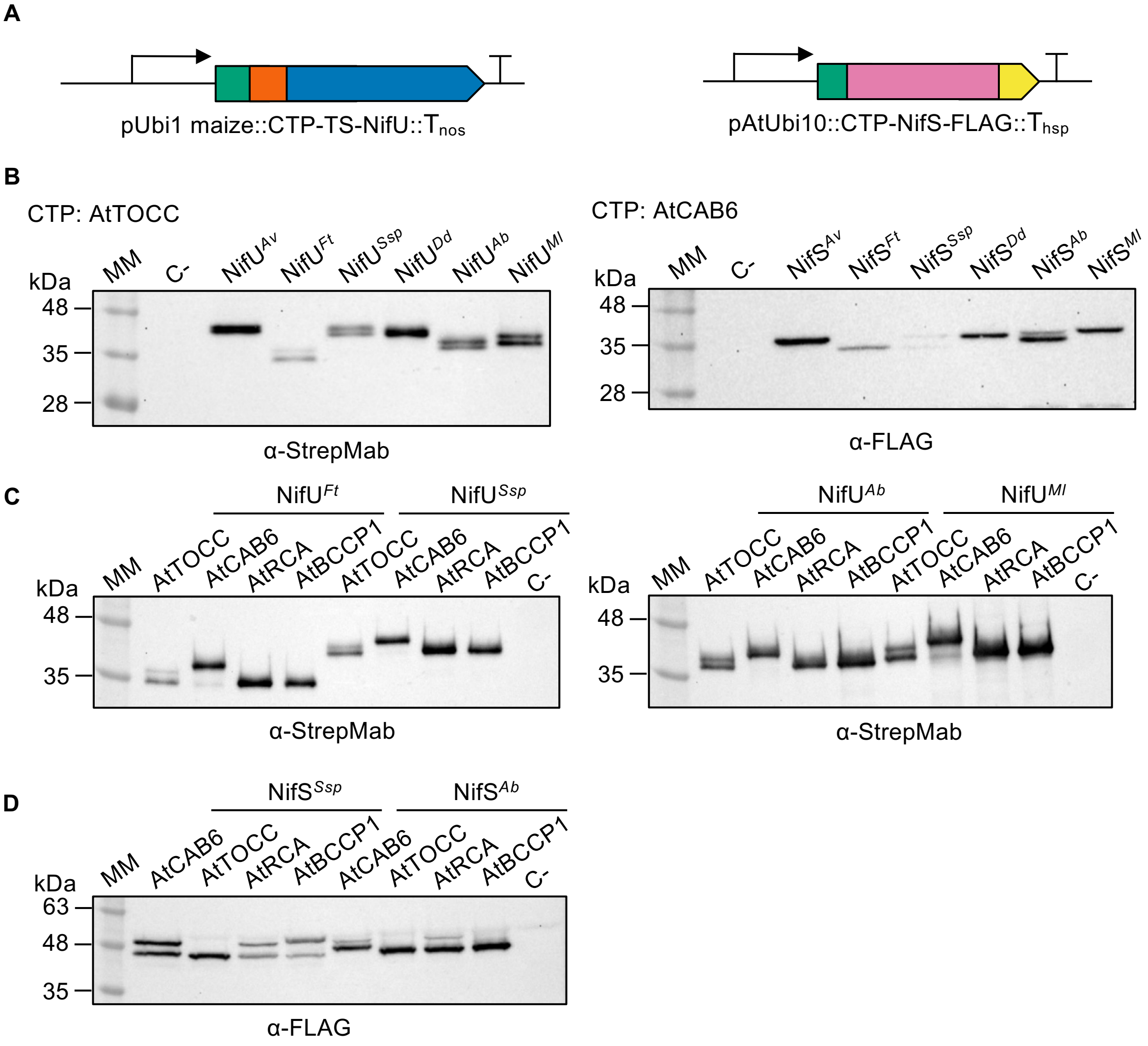


**Fig. S2.** Screening of chloroplast targeting peptides (CTP) to target NifU and NifS variants to rice chloroplasts. (A) Graphic representations of the genetic constructs showing the CTP (green), Twin-Strep tag (TS), orange, NifU (blue), NifS (pink), and FLAG tag (yellow). Initial (B) and refined rice protoplast CTP screening for NifU (C) and NifS (D) variants. Species abbreviations: *Azotobacter vinelandii* (*Av*), *Fischerella thermalis* (*Ft*), *Synechococcus sp.* JA-2-3B'a (*Ssp*), *Dechloromonas denitrificans* (*Dd*), *Azospirillum baldaniourum* (*Ab*), and *Marinobacterium lutimaris* (*Ml*). SDS-PAGE and immunoblot analysis performed with proteins extracts from protoplasts transformed with the *nifU* and *nifS* genes fused to different CTP (AtTOCC, AtCAB6, AtRCA and AtBCCP). Single bands suggest correct CTP processing. C- refers to protein extracts from non-transformed rice protoplasts.

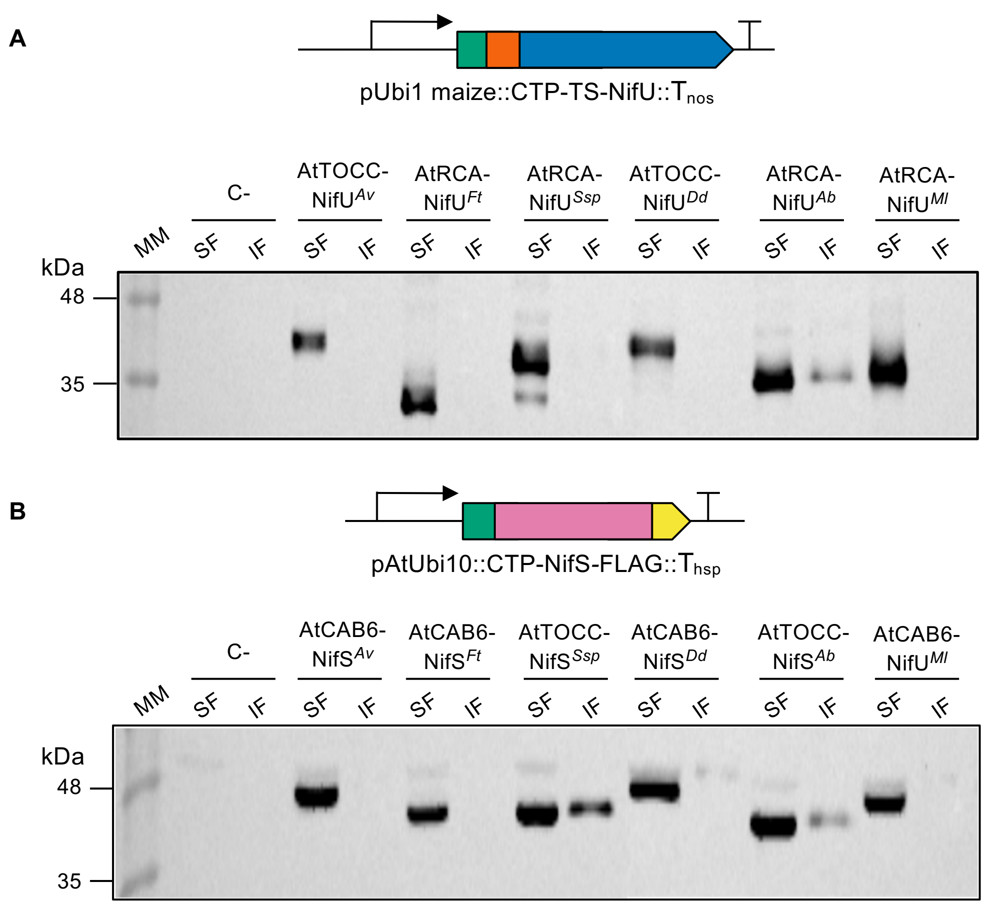


**Fig. S3.** Solubility screening of NifU (A) and NifS (B) variants with selected CTP in rice protoplasts. SDS-PAGE and immunoblot analysis performed with soluble (SF) and insoluble (IF) fractions from protoplasts transformed with the *nifU* and *nifS* genes fused to the CTP shown in Fig.1. Species abbreviations as described in Fig.1. C- refers to soluble and insoluble fractions from non-transformed rice protoplasts. Graphic representations of the genetic constructs are shown at the top of each panel: CTP (green), Twin-Strep tag (TS), orange, NifU (blue), NifS (pink), and FLAG tag (yellow).

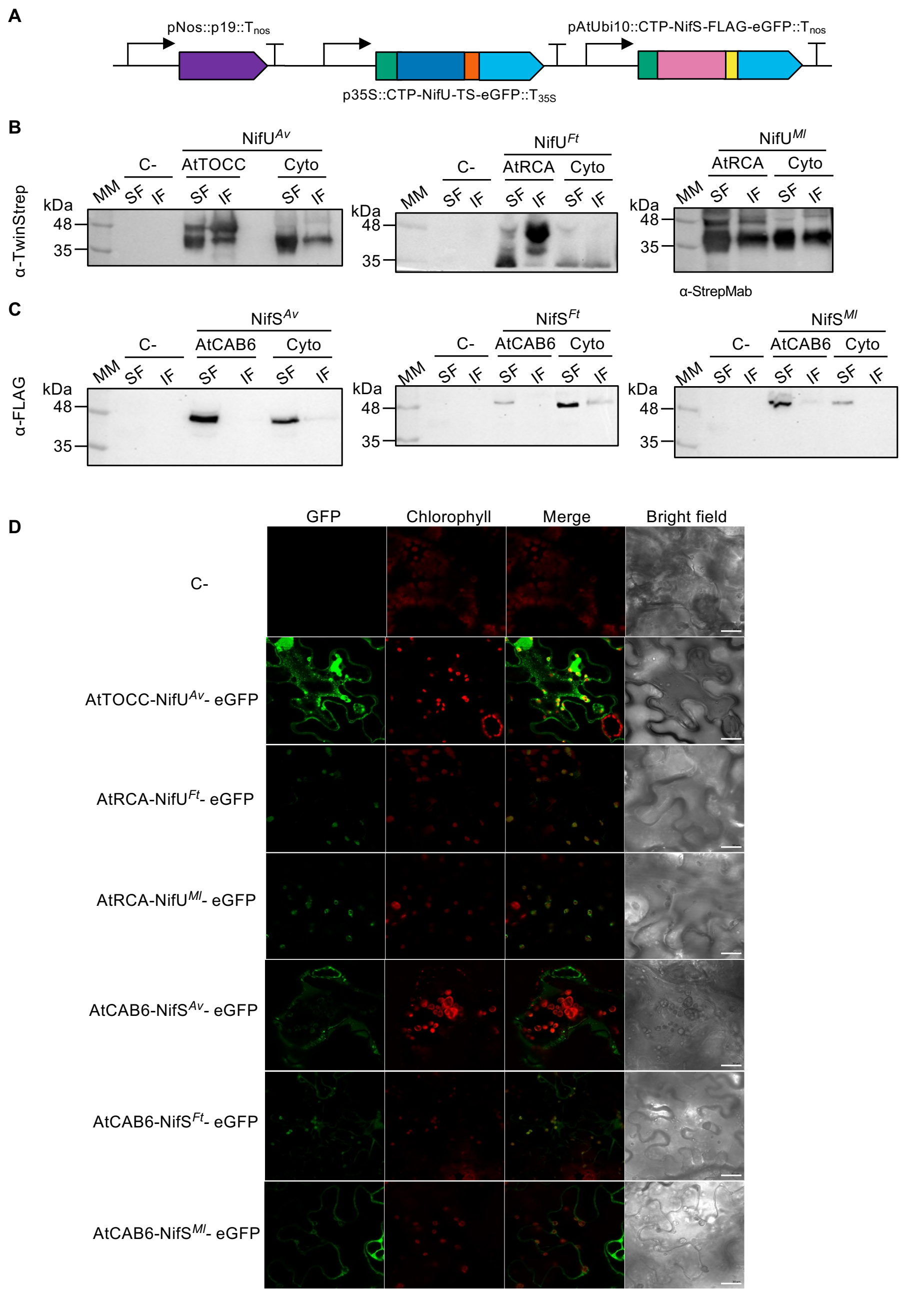


**Fig. S4.** Transient expression of selected NifU and NifS variants in *N. benthamiana* leaves. (A) Schematic representation of DNA constructs used to agroinfiltrate *N. benthamiana*. Graphic representations of the genetic constructs showing the CTP (green), Twin-Strep tag (TS), orange, NifU (blue), NifS (pink), FLAG tag (yellow), and eGFP (cyan). Constructs contained one NifUS pair from *A. vinelandii* (*Av*), *F. thermalis* (*Ft*), or *M. lutimaris* (*Ml*) and the p19 silencing suppressor. (B) SDS-PAGE and immunoblot analysis performed with soluble (SF) and insoluble (IF) fractions from with either the chloroplast-targeted or the cytosolic version of NifU and NifS. (C) Confocal microscopy images of the subcellular localization of NifU and NifS variants fused to a CTP and the green fluorescent protein (GFP) in agroinfiltrated *N. benthamiana*. Co-localization with chlorophyll autofluorescence (third row, merge) indicates correct import of NifU and NifS variants from *F. thermalis* and *M. lutimaris* into chloroplasts. Scale bar 10 µm. C-: non- transformed *N. benthamiana* leaves.

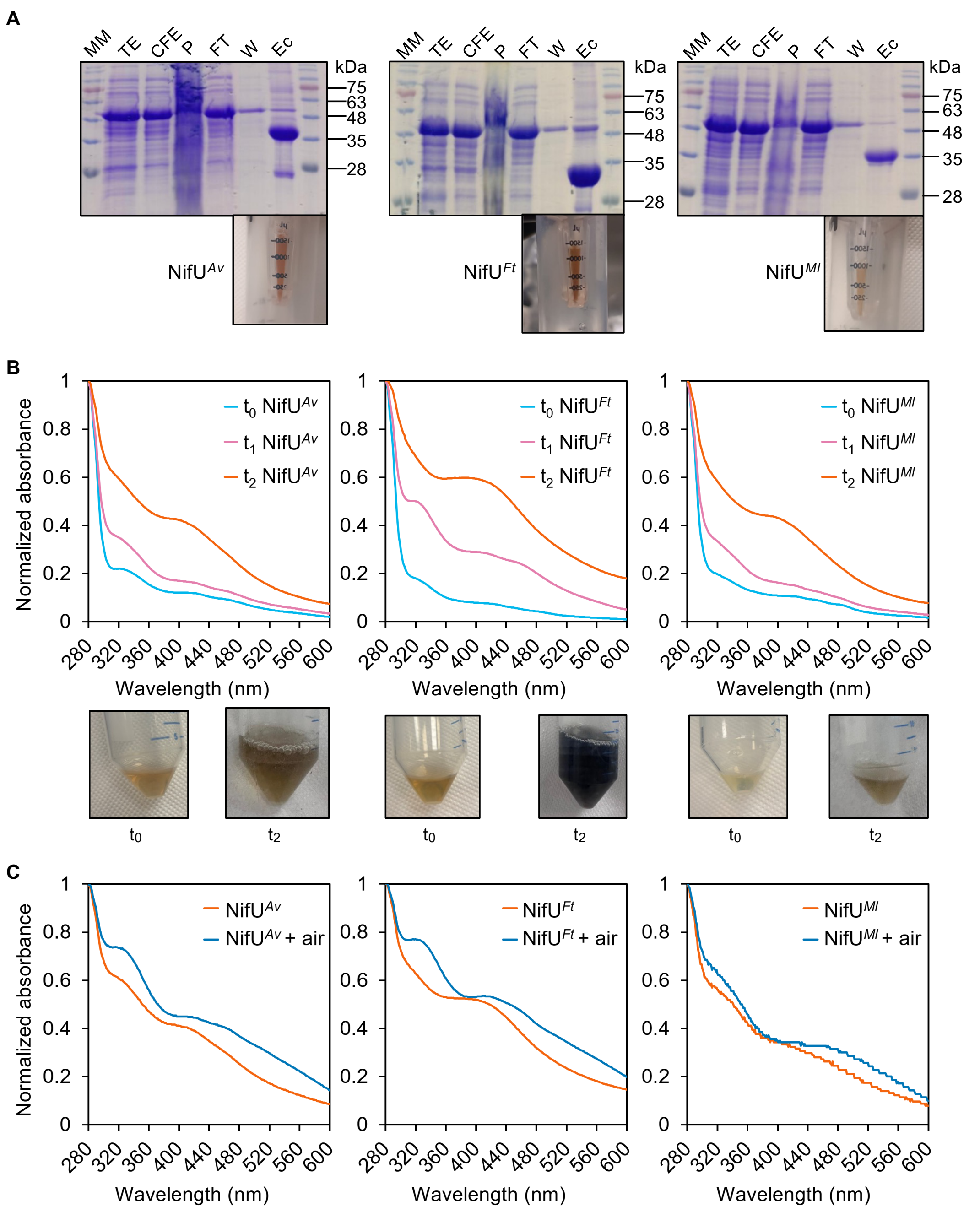


**Fig. S5.** Biochemical characterizations of NifU variants purified from *N. benthamiana* leaves. (A) Coomassie-stained SDS-PAGE analysis of the purification fractions from the *A. vinelandii* (NifU$^{Av}$, left), *F. thermalis* (NifU$^{Ft}$ middle), and *M. lutimaris* (NifU$^{Ml}$ right) strep-tagged NifU protein produced in *N. benthamiana*. Total Extract (TE), Cell Free Extract (CFE), Pellet (P), Flow-Through (FT), Wash (W) and Elution concentrated (Ec). A representative picture of each protein preparation is shown below the corresponding gel. (B) UV-vis spectra of [4Fe-4S] cluster reconstitution of NifU at t0 (before reconstitution, blue), t1 (3 h reconstitution, pink), and t2 (18h reconstitution, orange). NifU pictures at t0 and t2 are shown below each UV-vis panel. (C) UV-vis spectra of reconstituted NifU, before (orange) and after (blue) exposure to $O_2$ from air.

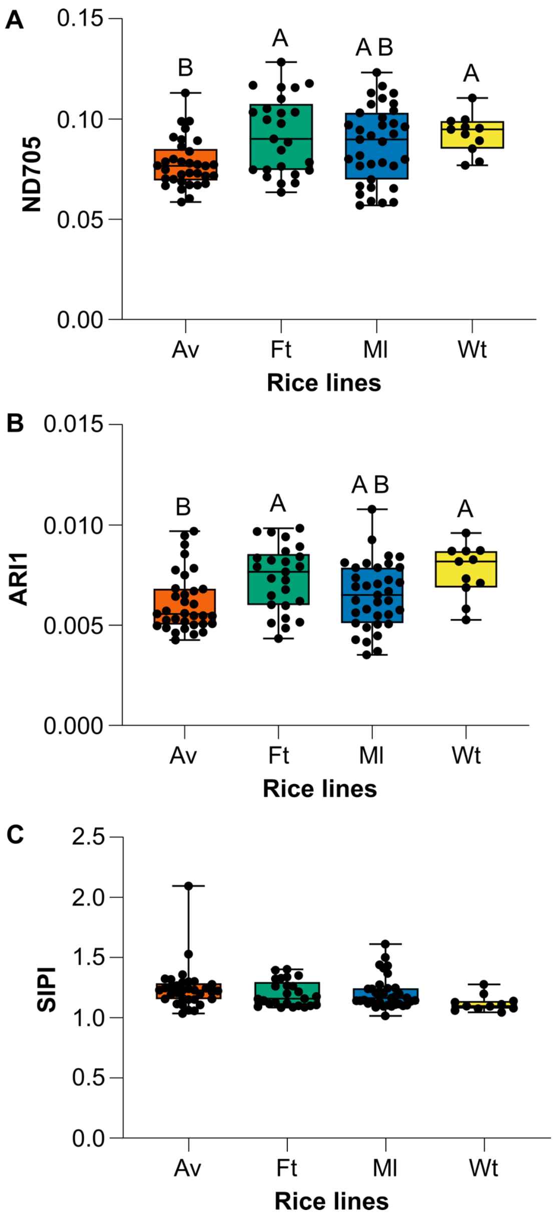


**Fig. S6.** Additional phenotype parameters of wild type (Wt) and transgenic *Ml*, *Ft*, and *Av* lines. Panels show boxplots in which each dot represents an individual plant, boxes indicate the interquartile range, and the central line represents the median. (A) Boxplots of Normalized Difference 705 Index (ND705) values. Statistical differences among phenotypes were assessed using the Kruskal–Wallis test followed by Dunn's multiple-comparison test. (B) Box plots of Anthocyanin Reflectance Index 1 (ARI1) values. Differences among phenotypes were assessed using a Type II analysis of variance (ANOVA), followed by Tukey–Kramer multiple-comparison tests. In (A) and (B), different letters above the boxes indicate significant differences between groups ($p < 0.05$), whereas groups sharing at least one letter are not significantly different. (C) Boxplots of Structure Insensitive Pigment Index (SIPI) values. Differences among phenotypes were assessed using a Type II analysis of variance (ANOVA) followed by Tukey–Kramer multiple-comparison tests. No significant differences among genotypes were detected ($p > 0.05$).

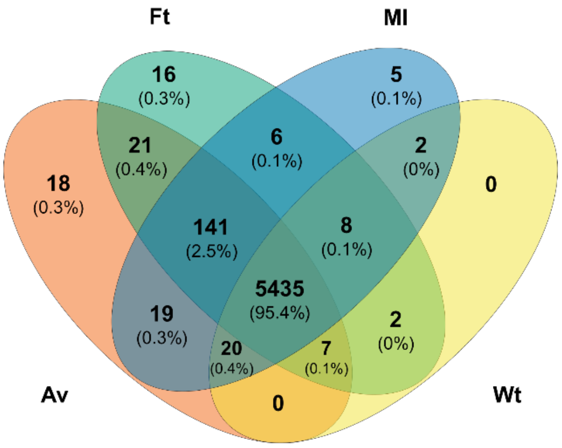


**Fig. S7**. Venn diagram showing the overlap of proteins identified in wild-type, *Av*, *Ft*, and *Ml* rice plants.

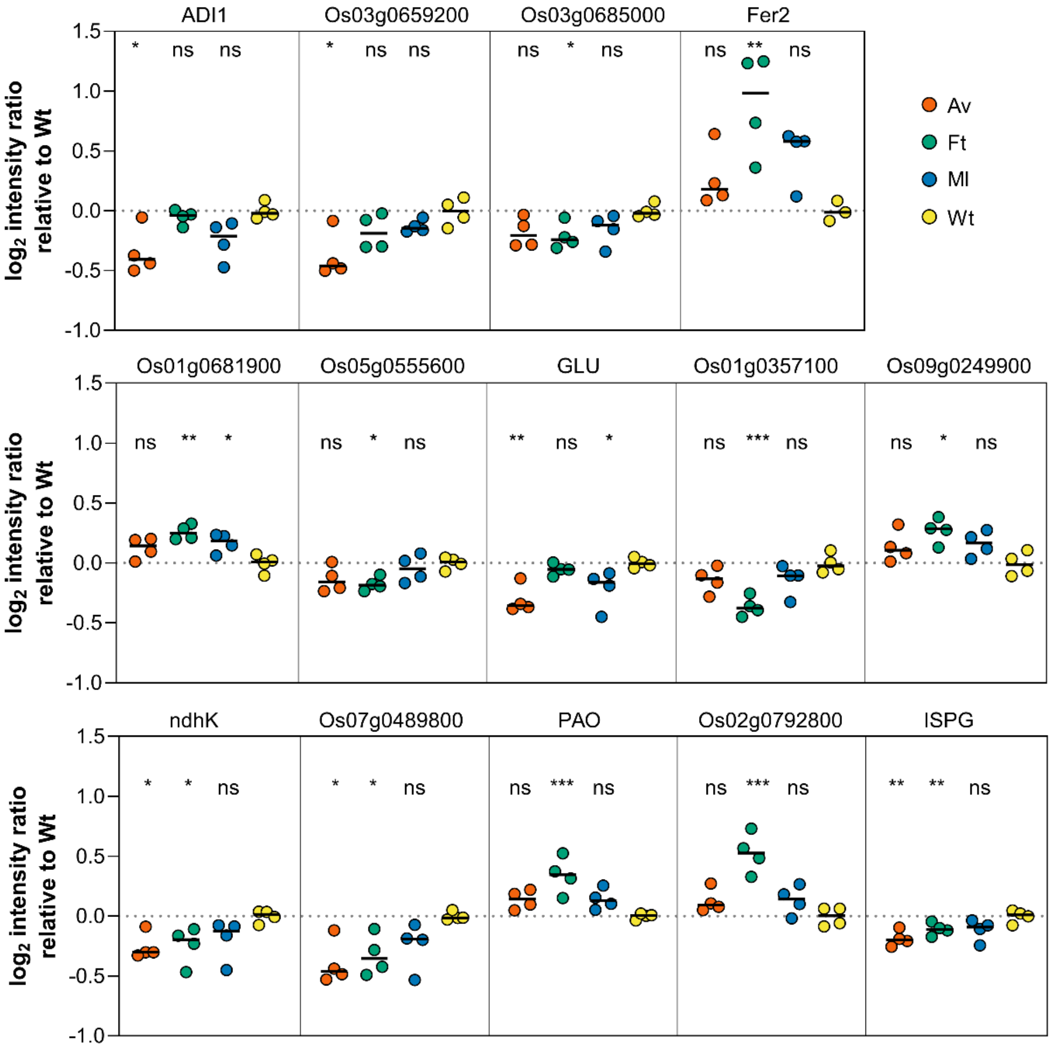


**Fig. S8.** Relative expression of [Fe-S] cluster proteins and ferritin in *Av*, *Ft* and *Ml* rice lines. Each dot represents the intensity value of each replicate after normalization to *wild-type* parametric mean, black bar indicates median. When normal distribution and homocedasticity was met, ONE-way ANOVA followed by test Dunnet test was done. Otherwise, Non-parametric ANOVA with Kruskal-Wallis test was performed. ns = $p < 0.05$, * = $p < 0.05$, ** = $p < 0.01$, *** = $p < 0.001$. Gene name (Protein name/description, UniProtKB): ADI1 (Ferredoxin-1, Q0J8M2), Os03g0659200 (Ferredoxin, fragment, A0A0P0W0Z6), Os03g0685000 (Ferredoxin, Q10F16), Fer2 (Ferritin, Q8L5K0), Os01g0681900 (Glutamate synthase 1 [NADH], Q0JKD0), Os05g0555600 (Glutamate synthase 2 [NADH], Q0DG35), GLU (Ferredoxin-dependent glutamate synthase, Q69RJ0) , Os01g0357100 (Ferredoxin nitrite reductase, chloroplastic; Ferredoxin nitrite reductase, Q0JMV6), Os09g0249900 (Ferredoxin thioredoxin reductase catalytic chain, Q6K471), *ndhK* (NAD(P)H-quinone oxidoreductase subunit K, P0C343), Os07g0489800 (Os07g0489800 protein, Q7XHS1), PAO (Pheophorbide a oxygenase, Q0DV66), Os02g0792800 (Os02g0792800 protein, Q6K689), ISPG (4-hydroxy-3-methylbut-2-en-1-yl diphosphate synthase (ferredoxin), Q6K8J4).

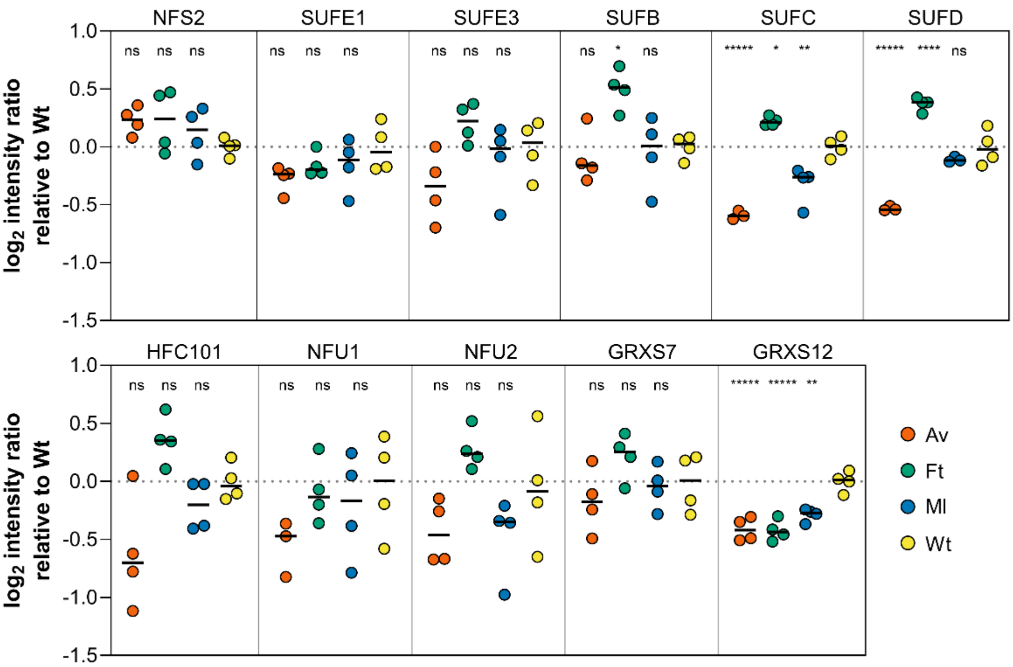


**Fig. S9.** Relative expression of [Fe-S] cluster biosynthesis proteins in *Av*, *Ft* and *Ml* rice lines. Each dot represents the intensity value of each replicate after normalization to *wild-type* parametric mean, black bar indicates median. Grubbs test was performed to detect outliers. When normal distribution and homocedasticity was met, ONE-way ANOVA followed by test Dunnet test was done. Otherwise, Non-parametric ANOVA with Kruskal-Wallis test was performed. ns = $p < 0.05$, * = $p < 0.05$, ** = $p < 0.01$, *** = $p < 0.001$, **** = $p < 0.0001$. Protein name ortholog from *A. thaliana* (Protein name/description, UniProtKB): NFS2 (cysteine desulfurase Fragment, Q0INV6) SUFE1 (Os09g0270900 protein Fragment, A0A0P0XJ88), SUFE3 (Quinolinate synthase, Q2QTL0), SUFB (Os01g0830000 protein, Q941V2), SUFC (ATP-dependent transporter ycf16, Q10LW5), SUFD (Os01g0127300 protein, Q5ZDZ8), HCF101 (Fe-S cluster assembly factor HCF101, Q0JJS8), NFU1 (NifU-like domain containing protein, Q10MC1). NFU2 (Nitrogen fixation protein, Q0IU70), GRXS7 (Monothiol glutaredoxin-S7, Q851Y7), GRXS12 (Monothiol glutaredoxin-S12, Q2QX01).

**Table S2**. G:Profiler pathway enrichment analysis of overexpressed proteins in *Av*, *Ft*, and *Ml* rice lines.

| Source | Term name | Term ID | $p_{adj}$ | Q | T∩Q | UniProtKB accession |
|---|---|---|---|---|---|---|
| GO:MF | Serine-type endopeptidase inhibitor activity | GO:0004867 | 5.02E-04 | 12 | 3 | Q0JR27, B7E9D7, A5HEI2 |
| | Endopeptidase inhibitor activity | GO:0004866 | 1.16E-03 | 12 | 3 | Q0JR27, B7E9D7, A5HEI2 |
| | Peptidase inhibitor activity | GO:0030414 | 1.16E-03 | 12 | 3 | Q0JR27, B7E9D7, A5HEI2 |
| | Endopeptidase regulator activity | GO:0061135 | 1.22E-03 | 12 | 3 | Q0JR27, B7E9D7, A5HEI2 |
| | Peptidase regulator activity | GO:0061134 | 1.27E-03 | 12 | 3 | Q0JR27, B7E9D7, A5HEI2 |
| | Enzyme inhibitor activity | GO:0004857 | 3.13E-02 | 12 | 3 | Q0JR27, B7E9D7, A5HEI2 |
| | Molecular function inhibitor activity | GO:0140678 | 3.18E-02 | 12 | 3 | Q0JR27, B7E9D7, A5HEI2 |
| GO:BP | Negative regulation of peptidase activity | GO:0010466 | 1.37E-03 | 12 | 3 | Q0JR27, B7E9D7, A5HEI2 |
| | Negative regulation of proteolysis | GO:0045861 | 1.37E-03 | 12 | 3 | Q0JR27, B7E9D7, A5HEI2 |
| | Regulation of peptidase activity | GO:0052547 | 1.37E-03 | 12 | 3 | Q0JR27, B7E9D7, A5HEI2 |
| | Negative regulation of hydrolase activity | GO:0051346 | 1.57E-03 | 12 | 3 | Q0JR27, B7E9D7, A5HEI2 |
| | Regulation of proteolysis | GO:0030162 | 1.68E-03 | 12 | 3 | Q0JR27, B7E9D7, A5HEI2 |
| | Regulation of hydrolase activity | GO:0051336 | 5.37E-03 | 12 | 3 | Q0JR27, B7E9D7, A5HEI2 |
| | Negative regulation of protein metabolic process | GO:0051248 | 2.00E-02 | 12 | 3 | Q0JR27, B7E9D7, A5HEI2 |
| | Negative regulation of molecular function | GO:0044092 | 2.06E-02 | 12 | 3 | Q0JR27, B7E9D7, A5HEI2 |
| | Negative regulation of catalytic activity | GO:0043086 | 2.06E-02 | 12 | 3 | Q0JR27, B7E9D7, A5HEI2 |
| | Galactose metabolic process | GO:0006012 | 2.54E-02 | 12 | 2 | Q8LNZ3, Q6YX79 |
| KEGG | Amino sugar and nucleotide sugar metabolism | KEGG:00520 | 1.14E-02 | 6 | 3 | O22080, Q8LNZ3, Q6YX79 |

Enrichment categories of upregulated proteins ($log_2$ FC > 1; q-value<0.05) present in transgenic *Av*, *Ft*, and *Ml* lines. Abbreviations: GO, Gene Ontology; MF, Molecular Function; BC, Biological Process; *Padj, p*-value adjusted for multiple testing using the g:SCS algorithm (*Padj* < 0.05); Q, number of upregulated proteins into Source category; T∩Q, number of overlapping proteins within the specific Term ID.

**Table S3.** Top 20 most upregulated or downregulated proteins in transgenic lines.

***Av* plants: upregulated**

| Accession | Gene Name | Protein Name | FC | p-value |
|---|---|---|---|---|
| Q306J3 | JAC1 | Dirigent protein | 54.3 | 0.00176 |
| Q75T45 | RSOsPR10 | Os12g0555000 protein | 30.0 | 0.00001 |
| O22080 | Os01g0660200 | chitinase | 24.5 | 0.00038 |
| Q2QNT0 | LOC_Os12g36850 | Os12g0555200 protein | 22.7 | 0.00139 |
| Q7F354 | Os01g0713200 | Os01g0713200 protein | 20.5 | 0.00064 |
| Q2QNN5 | LOC_Os12g37260 | Lipoxygenase | 18.5 | 0.00223 |
| Q10S66 | Cht11 | Chitinase 11 | 15.8 | 0.00174 |
| Q7XCK6 | Cht8 | Chitinase 8 | 14.8 | 0.00027 |
| Q5ZBR8 | Os01g0795000 | Os01g0795000 protein | 13.6 | 0.00044 |
| Q7F164 | Os01g0940700 | Os01g0940700 protein | 13.5 | 0.00082 |
| Q8L4V6 | LOC_Os10g38780 | Glutathione S-transferase | 13.4 | 0.00026 |
| Q8RU26 | Os01g0687400 | chitinase | 12.6 | 0.00027 |
| Q5VP99 | Os06g0582600 | Os06g0582600 protein | 12.3 | 0.00016 |
| Q8W2X5 | F3H-2 | Flavanone 3-dioxygenase 2 | 12.3 | 0.00019 |
| Q7Y0E8 | Os03g0405500 | Probable nucleoredoxin 1-1 | 10.8 | 0.00002 |
| Q0JR27 | Os01g0124100 | Os01g0124100 protein | 10.3 | 0.00027 |
| Q9FWU4 | LOC_Os10g34930 | Os10g0491000 protein | 9.7 | 0.00008 |
| Q93WY5 | Os09g0467200 | Glutathione transferase | 9.2 | 0.00007 |
| Q2QNS7 | LOC_Os12g36880 | Os12g0555500 protein | 8.5 | 0.00067 |
| Q7F1U0 | POX22.3 | Peroxidase 22.3 | 8.2 | 0.00026 |

***Av* plants: downregulated**

| Accession | Gene Name | Protein Name | FC | p-value |
|---|---|---|---|---|
| Q7XSN6 | Os04g0617900 | Germin-like protein 4-1 | 0.11 | 0.01083 |
| A0A0P0XHD8;Q6ZBP3 | Os08g0490900;H2B.2 | Histone H2B;Histone H2B.2 | 0.22 | 0.00008 |
| Q69T99 | AGPS1 | Glucose-1-phosphate adenylyltransferase small subunit 1, chloroplastic/amyloplastic | 0.27 | 0.00070 |
| A0A0P0V0F6 | Os01g0227800 | Os01g0227800 protein | 0.29 | 0.00747 |
| B7FA34 | Os05g0548900 | phosphoethanolamine N-methyltransferase | 0.33 | 0.01873 |
| Q2QVK7 | LOC_Os12g12470 | NADP-dependent oxidoreductase P1, putative, expressed | 0.34 | 0.00039 |
| A0A0P0WFX4 | Os04g0661600 | Molybdopterin biosynthesis protein CNX1 (Fragment) | 0.36 | 0.00012 |
| P0C422 | psbH | Photosystem II reaction center protein H | 0.36 | 0.02123 |

| | | | | |
|---|---|---|---|---|
| Q10N97 | LOC_Os03g16960 | 33 kDa secretory protein, putative, expressed | 0.36 | 0.04122 |
| Q851H6 | LOC_Os03g39830 | Expressed protein | 0.37 | 0.00063 |
| A0A0P0X302 | Os07g0164200 | Os07g0164200 protein | 0.40 | 0.00547 |
| Q7XHS1 | Os07g0489800 | Os07g0489800 protein | 0.41 | 0.00619 |
| Q10F62 | LOC_Os03g47610 | Os03g0679700 protein | 0.42 | 0.00364 |
| A0A0P0WL76 | Os05g0337400 | Os05g0337400 protein | 0.42 | 0.00011 |
| Q94EA4 | Os01g0731100 | Os01g0731100 protein | 0.42 | 0.02633 |
| Q10CL8 | LOC_Os03g54980 | Os03g0756800 protein | 0.43 | 0.00399 |
| Q0DKB2 | Os05g0177500 | Glycosyltransferase | 0.43 | 0.00057 |
| A0A0P0W0Z6;Q850T5 | Os03g0659200;LOC_Os03g45710 | Ferredoxin (Fragment);Ferredoxin | 0.43 | 0.01782 |
| Q6Z4N6 | R40G2 | Ricin B-like lectin R40G2 | 0.44 | 0.04553 |
| Q6EP57 | Os02g0578400 | Os02g0578400 protein | 0.44 | 0.00513 |

### *Ft* plants: upregulated

| Accession | Gene Name | Protein Name | FC | p-value |
|---|---|---|---|---|
| Q306J3 | JAC1 | Dirigent protein | 31.6 | 0.00235 |
| Q2QNN5 | LOC_Os12g37260 | Lipoxygenase | 29.1 | 0.00123 |
| Q7XCK6 | Cht8 | Chitinase 8 | 15.1 | 0.00002 |
| Q9FWU4 | LOC_Os10g34930 | Os10g0491000 protein | 13.9 | 0.00001 |
| Q10S66 | Cht11 | Chitinase 11 | 12.5 | 0.00177 |
| Q5ZBR8 | Os01g0795000 | Os01g0795000 protein | 12.5 | 0.00005 |
| Q8L5K0 | Fer2 | Ferritin | 12.5 | 0.00947 |
| O22080 | Os01g0660200 | chitinase | 11.8 | 0.00022 |
| Q8L4V6 | LOC_Os10g38780 | Glutathione S-transferase | 11.4 | 0.00006 |
| A0A0P0VQF9 | Os02g0783625 | Os02g0783625 protein | 11.2 | 0.00095 |
| Q7Y0E8 | Os03g0405500 | Probable nucleoredoxin 1-1 | 10.9 | 0.00007 |
| Q7F164 | Os01g0940700 | Os01g0940700 protein | 10.6 | 0.00021 |
| Q9LGK6 | Os01g0160800 | rRNA N-glycosylase | 10.4 | 0.00166 |
| Q7F354 | Os01g0713200 | Os01g0713200 protein | 10.1 | 0.00002 |
| Q5JMS4 | Os01g0962700 | Peroxidase | 9.0 | 0.00328 |
| Q6ZK52 | Os08g0127100 | Os08g0127100 protein | 8.8 | 0.02650 |
| Q10N21 | APX1 | L-ascorbate peroxidase 1, cytosolic | 8.3 | 0.00660 |
| Q93WY5 | Os09g0467200 | glutathione transferase | 8.3 | 0.00003 |
| Q2QNT0 | LOC_Os12g36850 | Os12g0555200 protein | 8.3 | 0.01534 |
| Q942C4 | Os01g0734800 | Glycosyltransferase | 8.2 | 0.00007 |

### *Ft* plants: downregulated

| Accession | Gene Name | Protein Name | FC | p-value |
|---|---|---|---|---|
| A3AHG5 | LEA17 | Late embryogenesis abundant protein 17 | 0.14 | 0.00061 |

| | | | | |
|---|---|---|---|---|
| Q2QVK7 | LOC_Os12g12470 | NADP-dependent oxidoreductase P1, putative, expressed | 0.22 | 0.01082 |
| A0A0P0XHD8; Q6ZBP3 | Os08g0490900;H2B.2 | Histone H2B;Histone H2B.2 | 0.23 | 0.00543 |
| Q69MM2 | Os09g0551600 | HMG type nucleosome/chromatin assembly factor | 0.25 | 0.01535 |
| Q53NL9 | LOC_Os11g47600 | Os11g0702100 protein | 0.29 | 0.00712 |
| A0A0P0XFE8 | Os08g0390200 | Os08g0390200 protein (Fragment) | 0.29 | 0.01700 |
| A0A0P0WN55;B7EYZ0 | Os05g0456300 | Os05g0456300 protein (Fragment);Os05g0456300 protein | 0.31 | 0.01447 |
| Q942A7 | Os01g0952100 | Germin-like protein 1-4 | 0.32 | 0.00259 |
| Q9FTN6 | Os01g0106300 | Os01g0106300 protein | 0.34 | 0.01806 |
| Q7XI46 | Os07g0622700 | Os07g0622700 protein | 0.34 | 0.00282 |
| Q5VME5 | CGTb | Glycosyltransferase | 0.35 | 0.02702 |
| Q84NP3 | OsFAD8 | FAD8 | 0.35 | 0.01483 |
| A0A0P0VJT6 | Os02g0530100 | Os02g0530100 protein | 0.35 | 0.00289 |
| Q0J845 | Os08g0137400 | Os08g0137400 protein | 0.37 | 0.00112 |
| Q6F382 | LOC_Os03g58100 | Os03g0795200 protein | 0.37 | 0.01164 |
| Q5SNH7 | Os01g0191100 | Os01g0191100 protein | 0.38 | 0.00892 |
| Q84M79 | LOC_Os03g64050 | Expressed protein | 0.38 | 0.01128 |
| Q7XXS4 | THI1 | Thiamine thiazole synthase, chloroplastic | 0.38 | 0.00138 |
| Q10SU0 | LOC_Os03g02040 | Os03g0111200 protein | 0.41 | 0.00233 |
| Q8S7N5 | LOC_Os10g42110 | non-specific serine/threonine protein kinase | 0.41 | 0.00689 |

### *Ml* plants: upregulated

| Accession | Gene Name | Protein Name | FC | p-value |
|---|---|---|---|---|
| O22080 | Os01g0660200 | chitinase | 5.6 | 0.0014 |
| A0A0P0UXQ2 | Os01g0139000 | Reticulon-like protein | 5.5 | 0.0011 |
| Q7XS61 | Os04g0165400 | Os04g0165400 protein | 5.1 | 0.0034 |
| A0A0P0VQF9 | Os02g0783625 | Os02g0783625 protein | 4.1 | 0.0038 |
| Q5JMS0 | Os01g0963300 | Os01g0963300 protein | 3.7 | 0.0012 |
| Q0JR27 | Os01g0124100 | Os01g0124100 protein | 3.7 | 0.0037 |
| Q7Y0E8 | Os03g0405500 | Probable nucleoredoxin 1-1 | 3.6 | 0.0022 |
| Q6I5I7 | Os05g0467300 | Os05g0467300 protein | 3.3 | 0.0025 |
| Q8LHL0 | Os07g0623300 | Os07g0623300 protein | 3.2 | 0.0002 |
| Q2QRV3 | PIOX | Alpha-dioxygenase PIOX | 3.0 | 0.0003 |
| Q69IN8 | Os09g0513000 | Os09g0513000 protein | 3.0 | 0.0013 |

| | | | | |
|---|---|---|---|---|
| P0C2Y7 | atpI | ATP synthase subunit a, chloroplastic | 2.9 | 0.0013 |
| Q942C4 | Os01g0734800 | Glycosyltransferase | 2.8 | 0.0044 |
| Q10Q18 | LOC_Os03g11440 | Os03g0213100 protein | 2.8 | 0.0005 |
| B7E9D7 | Os01g0124650 | Os01g0124650 protein | 2.4 | 0.0042 |
| Q8LNZ3 | UGE-1 | UDP-glucose 4-epimerase 1 | 2.4 | 0.0011 |
| A0A0P0XQB2 | Os09g0557800 | Transmembrane 9 superfamily member (Fragment) | 2.3 | 0.0013 |
| Q0DRZ2;Q10LL5 | Os03g0344100;LOC_Os03g22380 | Os03g0344100 protein | 2.3 | 0.0012 |
| A5HEI2 | pinA | Bowman-Birk type proteinase inhibitor A | 2.2 | 0.0014 |
| Q6EPN8 | FACE1 | CAAX prenyl protease 1 homolog | 2.2 | 0.0005 |

#### ***Ml*** **plants: downregulated**

| **Accession** | **Gene Name** | **Protein Name** | **FC** | **p-value** |
|---|---|---|---|---|
| Q851H6 | LOC_Os03g39830 | Expressed protein | 0.44 | 0.0035 |
| A0A0P0V0F6 | Os01g0227800 | Os01g0227800 protein | 0.47 | 0.0001 |

FC = fold-change in lineal scale; q-value < 0.05.

**Table S4.** G:Profiler pathway enrichment analysis of downregulated proteins in *Av* rice lines (but not in *Ft*, or *Ml* rice lines).

| Source | Term name | Term ID | $p_{adj}$ | Q | T∩Q | UniProtKB accession |
|---|---|---|---|---|---|---|
| GO:MF | Catalytic activity | GO:0003824 | 1.24E-03 | 111 | 83 | Q7F354, Q2QNN5, Q10S66, Q7XCK6, Q5ZBR8, Q7F164, Q8L4V6, Q8RU26, Q5VP99, Q8W2X5, Q93WY5, Q7F1U0, Q7XKV4, Q7XEL9, Q42993, A0A0P0XA49, Q0DCP0, A0A0P0XUN5, Q6EQK1, Q0JLY2, Q8GS76, Q6Z674, Q2QVJ8, Q7XL00, Q0DIB3, Q2QXB3, Q5JMS4, Q9LGK6, Q0JCV5, Q5Z5T3, Q6Z0T7, Q941Y8, Q2QN11, Q945W6, Q9AS12, Q6ZGM5, Q8H620, Q5ZCF2, A0A0P0WU39, Q0JK51, A3AQC6, Q338Z5, Q2R5M2, Q5Z7J2, Q6ZIH1, Q0DZJ2, Q5Z7I5, Q6K6I2, Q6Z481, Q10P89, Q5JNC0, Q53NE7, Q0ILB9, Q0DU12, Q5WMR4, A0A0P0VFA8, P42211, Q84LM2, Q10PS7, Q6K7A3, Q5VP37, Q5W6C5, Q9AS85, Q65XC3, Q7XSU8, A0A0P0WZL7, Q10N20, Q7X8G7, Q7Y0F2, Q657L8, Q10RH3, Q0D5S1, Q8S0I1, A0A0P0X0W6, Q9S827, Q6YZ09, Q5W745, Q84S65, Q8S1G9, Q06398, A0A0P0V2Z4, A0A0P0V9F2, Q69P84 |
| | Oxidoreductase activity | GO:0016491 | 1.94E-04 | 111 | 28 | Q2QNN5, Q8W2X5, Q7F1U0, A0A0P0XA49, Q0DCP0, Q2QVJ8, Q7XL00, Q2QXB3, Q5JMS4, Q0JCV5, Q5Z5T3, Q941Y8, Q9AS12, Q6ZGM5, Q8H620, A0A0P0WU39, Q5Z7J2, Q6ZIH1, Q5Z7I5, Q10PS7, Q7XSU8, Q7X8G7, Q7Y0F2, Q9S827, Q6YZ09, Q5W745, Q8S1G9, Q69P84 |
| | Peptidase activity | GO:0008233 | 4.02E-03 | 111 | 15 | Q5ZBR8, Q5VP99, Q6Z674, Q0DIB3, Q2QN11, Q5ZCF2, Q2R5M2, Q6K6I2, Q10P89, Q53NE7, P42211, Q84LM2, Q6K7A3, Q5W6C5, A0A0P0V9F2 |
| | Endopeptidase activity | GO:0004175 | 1.94E-03 | 111 | 12 | Q5ZBR8, Q5VP99, Q6Z674, Q0DIB3, Q2QN11, Q5ZCF2, Q6K6I2, Q10P89, Q53NE7, P42211, Q84LM2, Q6K7A3 |
| | Hydrolase activity, acting on glycosyl bonds | GO:0016798 | 1.23E-02 | 111 | 12 | Q7F354, Q10S66, Q7XCK6, Q7F164, Q8RU26, Q7XKV4, Q7XEL9, Q42993, A0A0P0XUN5, Q9LGK6, Q9AS85, Q0D5S1 |
| | Antioxidant activity | GO:0016209 | 1.26E-02 | 111 | 8 | Q7F1U0, A0A0P0XA49, Q0DCP0, Q5JMS4, Q9AS12, Q5Z7J2, Q7XSU8, Q7Y0F2 |
| | Lactoperoxidase activity | GO:0140825 | 5.04E-03 | 111 | 7 | Q7F1U0, A0A0P0XA49, Q0DCP0, Q5JMS4, Q9AS12, Q5Z7J2, Q7XSU8 |
| | Peroxidase activity | GO:0004601 | 4.13E-02 | 111 | 7 | Q7F1U0, A0A0P0XA49, Q0DCP0, Q5JMS4, Q9AS12, Q5Z7J2, Q7XSU8 |
| | Oxidoreductase activity, acting on peroxide as acceptor | GO:0016684 | 4.28E-02 | 111 | 7 | Q7F1U0, A0A0P0XA49, Q0DCP0, Q5JMS4, Q9AS12, Q5Z7J2, Q7XSU8 |
| | Aspartic-type peptidase activity | GO:0070001 | 4.00E-02 | 111 | 6 | Q6Z674, Q0DIB3, Q2QN11, Q6K6I2, Q53NE7, P42211 |
| | Aspartic-type endopeptidase activity | GO:0004190 | 4.00E-02 | 111 | 6 | Q6Z674, Q0DIB3, Q2QN11, Q6K6I2, Q53NE7, P42211 |
| | Chitinase activity | GO:0004568 | 2.88E-03 | 111 | 5 | Q10S66, Q7XCK6, Q8RU26, Q42993, A0A0P0XUN5 |
| | NADH dehydrogenase activity | GO:0003954 | 6.61E-03 | 111 | 3 | Q941Y8, Q6YZ09, Q5W745 |
| GO:BP | Response to chemical | GO:0042221 | 2.67E-04 | 95 | 24 | Q75T45, Q2QNT0, Q8L4V6, Q93WY5, Q2QNS7, Q7F1U0, A0A0P0XA49, Q0DCP0, Q0JLY2, |

| | | | | | | |
|---|---|---|---|---|---|---|
| GO:BP | | | | | | Q5JMS4, Q10PW9, Q945W6, Q9AS12, Q5Z7J2, Q69RN2, Q0ILB9, Q9SXF8, Q7XSU8, Q10N20, Q7Y0F2, A0A0P0WDN1, Q6YYB0, Q06398, Q69P84 |
| | Proteolysis | GO:0006508 | 1.88E-05 | 95 | 17 | Q5ZBR8, Q5VP99, Q6Z674, Q0DIB3, Q2QN11, Q5ZCF2, Q2R5M2, Q6K6I2, Q10P89, Q53NE7, P42211, Q0JR29, Q84LM2, Q6K7A3, Q5W6C5, Q10J94, A0A0P0V9F2 |
| | Response to toxic substance | GO:0009636 | 2.91E-03 | 95 | 10 | Q7F1U0, A0A0P0XA49, Q0DCP0, Q5JMS4, Q9AS12, Q5Z7J2, Q0ILB9, Q7XSU8, Q7Y0F2, A0A0P0WDN1 |
| | Cellular oxidant detoxification | GO:0098869 | 1.78E-03 | 95 | 9 | Q7F1U0, A0A0P0XA49, Q0DCP0, Q5JMS4, Q9AS12, Q5Z7J2, Q7XSU8, Q7Y0F2, A0A0P0WDN1 |
| | Cellular detoxification | GO:1990748 | 3.55E-03 | 95 | 9 | Q7F1U0, A0A0P0XA49, Q0DCP0, Q5JMS4, Q9AS12, Q5Z7J2, Q7XSU8, Q7Y0F2, A0A0P0WDN1 |
| | Cellular response to toxic substance | GO:0097237 | 3.68E-03 | 95 | 9 | Q7F1U0, A0A0P0XA49, Q0DCP0, Q5JMS4, Q9AS12, Q5Z7J2, Q7XSU8, Q7Y0F2, A0A0P0WDN1 |
| | Detoxification | GO:0098754 | 1.60E-02 | 95 | 9 | Q7F1U0, A0A0P0XA49, Q0DCP0, Q5JMS4, Q9AS12, Q5Z7J2, Q7XSU8, Q7Y0F2, A0A0P0WDN1 |
| | Reactive oxygen species metabolic process | GO:0072593 | 2.76E-03 | 95 | 8 | Q7F1U0, A0A0P0XA49, Q0DCP0, Q5JMS4, Q9AS12, Q5Z7J2, Q7XSU8, A0A0P0WDN1 |
| | Hydrogen peroxide catabolic process | GO:0042744 | 9.18E-03 | 95 | 7 | Q7F1U0, A0A0P0XA49, Q0DCP0, Q5JMS4, Q9AS12, Q5Z7J2, Q7XSU8 |
| | Hydrogen peroxide metabolic process | GO:0042743 | 1.11E-02 | 95 | 7 | Q7F1U0, A0A0P0XA49, Q0DCP0, Q5JMS4, Q9AS12, Q5Z7J2, Q7XSU8 |
| GO:CC | Extracellular region | GO:0005576 | 1.44E-06 | 89 | 19 | Q306J3, Q5ZBR8, Q8RU26, Q5VP99, Q7F1U0, Q7XKV4, A0A0P0XA49, Q0DCP0, Q5JMS4, Q5Z5T3, Q2QN11, Q9AS12, Q5Z7J2, Q0JR29, Q6K7A3, Q7XSU8, Q10J94, Q6ESF0, A0A0P0V9F2 |
| KEGG | Metabolic pathways | KEGG:01100 | 1.96E-02 | 38 | 28 | Q2QNN5, Q10S66, Q7XCK6, Q8L4V6, Q8RU26, Q93WY5, Q7XKV4, A0A0P0XA49, Q8H4P7, Q0JLY2, Q5JMS4, Q5Z5T3, Q6Z0T7, Q9AS12, A0A0P0WU39, A3AQC6, Q5JNC0, Q0ILB9, A0A0P0VFA8, Q7XSU8, Q7X8G7, Q8S0I1, Q9S827, Q5W745, Q8S1G9, Q06398, A0A0P0V2Z4, Q69P84 |

Enrichment categories of upregulated proteins ($log_2$ FC > 1; q-value<0.05). Abbreviations: GO, Gene Ontology; MF, Molecular Function; BC, Biological Process; *Padj, p*-value adjusted for multiple testing using the g:SCS algorithm (*Padj* < 0.05); Q, number of upregulated proteins into Source category; T∩Q, number of overlapping proteins within the specific Term ID.

**Table S5**. G:Profiler pathway enrichment analysis of overexpressed proteins in *Av* and *Ft* rice lines (but not in *Ml* lines).

| Source | Term name | Term ID | $p_{adj}$ | Q | T∩Q | UniProtKB accession |
|---|---|---|---|---|---|---|
| GO:MF | Hydrolase activity | GO:0016787 | 1.17E-03 | 121 | 42 | Q0J0N4, Q6EPR6, Q0JF91, Q6YVU7, A0A0P0V5L9, Q0J105, Q6L4X7, A0A0P0WQZ8, Q6K6B3, Q5JLW5, Q10QV1, A3C4N5, Q10R90, A0A0P0XKC7, Q94ED2, Q651M0, Q75K72, B7EIC7, Q10QA5, Q0IU77, Q8LHP5, A0A0P0VUV1, Q5WMX0, Q0JL73, Q7F1K9, Q337C4, Q9LGA5, Q2R1Y6, Q7XIV4, Q10PD0, Q6YVU4, Q0JQZ2, Q84NP7, P51823, Q06396, O49827, A0A0P0Y5Y3, Q75LI0, A0A0P0VFX7, Q9LGZ3, A0A0P0YBQ1, Q60EH3 |
| | Hydrolase activity, hydrolyzing O-glycosyl compounds | GO:0004553 | 7.19E-03 | 121 | 12 | Q0J0N4, Q6YVU7, Q94ED2, Q75K72, Q5WMX0, Q7XIV4, Q6YVU4, Q0JQZ2, O49827, Q75LI0, A0A0P0VFX7, Q60EH3 |
| | Glucosidase activity | GO:0015926 | 3.01E-02 | 121 | 6 | Q0J0N4, Q6YVU7, Q94ED2, Q6YVU4, Q0JQZ2, Q60EH3 |
| | Hydrolase activity, acting on glycosyl bonds | GO:0016798 | 1.51E-02 | 121 | 12 | Q0J0N4, Q6YVU7, Q94ED2, Q75K72, Q5WMX0, Q7XIV4, Q6YVU4, Q0JQZ2, O49827, Q75LI0, A0A0P0VFX7, Q60EH3 |
| GO:CC | Precatalytic spliceosome | GO:0071011 | 2.97E-02 | 114 | 4 | Q10PA9, Q6YU78, Q5WMV0, Q0J7Y5 |
| | pICln-Sm protein complex | GO:0034715 | 4.12E-02 | 114 | 3 | Q10PA9, Q6YU78, Q5WMV0 |
| | Spliceosomal tri-snRNP complex | GO:0097526 | 4.55E-02 | 114 | 4 | Q10PA9, Q6YU78, Q5WMV0, Q0J7Y5 |
| | U5 snRNP | GO:0005682 | 4.23E-03 | 114 | 3 | Q10PA9, Q6YU78, Q5WMV0 |
| | Endomembrane system | GO:0012505 | 9.56E-03 | 114 | 24 | Q10EM0, Q2QPH4, Q8S2K0, Q9FTT3, Q10AZ7, Q2QWZ9, Q53PC7, Q8LHP5, Q851A2, Q7F1K9, Q5W6W7, Q2R1Y6, Q7XI39, P51823, Q06396, Q0DJC5, Q69SG3, Q6YU98, Q2QNU0, Q75GB3, Q0IT04, Q10CF7, A0A0P0W2T6, Q6YZD2 |
| | Coated vesicle | GO:0030135 | 4.76E-03 | 114 | 6 | Q53PC7, Q851A2, Q7XI39, Q0IT04, Q10CF7, Q6YZD2 |
| | U1 snRNP | GO:0005685 | 1.76E-02 | 114 | 3 | Q10PA9, Q6YU78, Q5WMV0 |
| | COPII-coated ER to Golgi transport vesicle | GO:0030134 | 4.30E-02 | 114 | 4 | Q851A2, Q0IT04, Q10CF7, Q6YZD2 |
| | U2 snRNP | GO:0005686 | 5.00E-02 | 114 | 3 | Q10PA9, Q6YU78, Q5WMV0 |
| KEGG | Fatty acid degradation | KEGG:00071 | 3.59E-03 | 45 | 5 | Q2R1G8, Q10QN9, A0A0P0V6M8, Q5W6W7, Q2R8Z5 |

Enrichment categories of exclusively upregulated proteins (log2 FC > 1; q-value<0.05). Abbreviations: GO, Gene Ontology; MF, Molecular Function; BC, Biological Process; *Padj, p*-value adjusted for multiple testing using the g:SCS algorithm (*Padj* < 0.05); Q, number of upregulated proteins into Source category; T∩Q, number of overlapping proteins within the specific Term ID.

**Table S6.** Proteins detected only in specific *Av*, *Ft* or *Ml* rice lines.

| Transgenic plant | Accession | Gene Name | Protein Name |
|---|---|---|---|
| *Av* | A0A0P0V2C2 | Os01g0326300 | Peroxidase (Fragment) |
| | A0A0P0VED8 | Os02g0131100 | Os02g0131100 protein (Fragment) |
| | A0A0P0W7L8 | Os04g0178300; CPS4 | syn-copalyl-diphosphate synthase (Fragment);Syn-copalyl diphosphate synthase, chloroplastic |
| | A0A0P0Y5F2 | Os11g0687200;LOC_Os11g46000 | Os11g0687200 protein;Expressed protein |
| | B7FAK3 | Os04g0577800 | Os04g0577800 protein |
| | Q0D5S8 | Os07g0538400 | Os07g0538400 protein (Fragment) |
| | Q0D879 | Os07g0178800 | Os07g0178800 protein |
| | Q0J3Z5 | Os08g0544500;NAP1 | Os08g0544500 protein (Fragment);Probable protein NAP1 |
| | Q0JAF0 | Os04g0602500 | Pectin acetylesterase |
| | Q10MX3 | Os03g0291500 | Asparagine synthetase [glutamine-hydrolyzing] 1 |
| | Q2QZH3 | LOC_Os11g45990 | Os11g0687100 protein |
| | Q5JKV6 | Os01g0742400 | non-specific serine/threonine protein kinase |
| | Q69IV0 | P0498F03.1 | Glycosyltransferase |
| | Q7FAE1 | Os04g0179200 | Momilactone A synthase |
| | Q8LQ36 | Os01g0851400 | Putative ataxin-3 homolog |
| | Q9LHU2 | Os01g0222600 | Os01g0222600 protein |
| | U00001 | nifU_Av | NifU_Av |
| | U00002 | nifS_Av | NifS_Av |
| *Ft* | A0A0P0VPM6 | Os02g0755900 | Glycosyltransferase (Fragment) |
| | A0A0P0XN78 | Os09g0448200;HAK17 | Probable potassium transporter 17 |
| | Q0DBM6 | Os06g0549900 | Os06g0549900 protein |
| | Q0DLF7 | Os05g0104200 | Os05g0104200 protein |
| | Q10MS8 | LOC_Os03g18550 | Mitochondrial carrier protein, expressed |
| | Q2QUC5 | CYP71P1 | Tryptamine 5-hydroxylase |
| | Q5ZAF2 | Os01g0597800 | Glycosyltransferase |
| | Q6ER51 | prx30 | Peroxidase |
| | Q6H545 | IPK2 | Inositol polyphosphate multikinase IPK2 |
| | Q6Z564 | Os08g0412700 | Os08g0412700 protein |
| | Q7XIW9 | prx115 | Peroxidase |
| | Q7XWU3 | CAD6 | Probable cinnamyl alcohol dehydrogenase 6 |
| | Q8RUJ2 | LOC_Os10g38340 | glutathione transferase |
| | U00003 | nifU_Ft | NifU_Ft |
| | U00004 | nifS_Ft | NifS_Ft |
| | U00008 | hptII | hptII |
| *Ml* | A0A0P0W2P0 | Os03g0736300;GLU2 | Endoglucanase; Endoglucanase 10 |
| | Q2QYK6 | LOC_Os12g02370 | Chalcone-flavonone isomerase family protein |
| | Q6AUC6 | QSOX1 | Sulfhydryl oxidase 1 |
| | U00005 | nifU_Ml | NifU_Ml |
| | U00006 | nifS_Ml | NifS_Ml |

**Table S7**. G:Profiler pathway enrichment analysis of downregulated proteins in *Av* rice lines (but not in *Ft* or *Ml* lines).

| Source | Term name | Term ID | $p_{adj}$ | Q | T∩Q | UniProtKB accession |
|---|---|---|---|---|---|---|
| GO:MF | Metal cluster binding | GO:0051540 | 7.63E-03 | 23 | 4 | Q10F62, A0A0P0W0Z6, Q0J8M2, Q69RJ0 |
| | Iron-sulfur cluster binding | GO:0051536 | 7.63E-03 | 23 | 4 | Q10F62, A0A0P0W0Z6, Q0J8M2, Q69RJ0 |
| GO:BP | Generation of precursor metabolites and energy | GO:0006091 | 5.80E-04 | 22 | 7 | Q69T99, A0A0P0W0Z6, Q6EP57, Q0J8M2, Q0D5P8, Q84R40, Q6ZDP0 |
| | Photosynthesis | GO:0015979 | 8.08E-05 | 22 | 6 | P0C422, Q6EP57, Q0D5P8, Q84R40, Q6ZDP0, Q6Z8N7 |
| | Electron transport chain | GO:0022900 | 9.96E-05 | 22 | 6 | A0A0P0W0Z6, Q6EP57, Q0J8M2, Q0D5P8, Q84R40, Q6ZDP0 |
| | Photosynthetic electron transport chain | GO:0009767 | 7.35E-05 | 22 | 4 | Q6EP57, Q0D5P8, Q84R40, Q6ZDP0 |
| | Photosynthesis, light reaction | GO:0019684 | 2.26E-03 | 22 | 4 | Q6EP57, Q0D5P8, Q84R40, Q6ZDP0 |
| GO:CC | Chloroplast | GO:0009507 | 5.11E-08 | 25 | 13 | Q69T99, P0C422, Q10F62, A0A0P0W0Z6, Q6EP57, Q6ATB2, Q0J8M2, Q0D5P8, Q7FB12, A0A0P0Y0Q1, P0C587, Q69RJ0, Q69SV0 |
| | Plastid | GO:0009536 | 1.17E-07 | 25 | 13 | Q69T99, P0C422, Q10F62, A0A0P0W0Z6, Q6EP57, Q6ATB2, Q0J8M2, Q0D5P8, Q7FB12, A0A0P0Y0Q1, P0C587, Q69RJ0, Q69SV0 |
| | Thylakoid | GO:0009579 | 9.39E-08 | 25 | 8 | P0C422, Q6EP57, Q0D5P8, Q7FB12, A0A0P0Y0Q1, B9FBM2, Q69SV0, Q6Z8N7 |
| | Thylakoid membrane | GO:0042651 | 5.38E-07 | 25 | 7 | P0C422, Q6EP57, Q0D5P8, Q7FB12, A0A0P0Y0Q1, Q69SV0, Q6Z8N7 |
| | Photosynthetic membrane | GO:0034357 | 9.31E-07 | 25 | 7 | P0C422, Q6EP57, Q0D5P8, Q7FB12, A0A0P0Y0Q1, Q69SV0, Q6Z8N7 |
| | Chloroplast thylakoid membrane | GO:0009535 | 2.12E-04 | 25 | 5 | P0C422, Q0D5P8, Q7FB12, A0A0P0Y0Q1, Q69SV0 |
| | Plastid thylakoid membrane | GO:0055035 | 2.12E-04 | 25 | 5 | P0C422, Q0D5P8, Q7FB12, A0A0P0Y0Q1, Q69SV0 |
| | Plastid thylakoid | GO:0031976 | 4.75E-04 | 25 | 5 | P0C422, Q0D5P8, Q7FB12, A0A0P0Y0Q1, Q69SV0 |
| | Chloroplast thylakoid | GO:0009534 | 4.75E-04 | 25 | 5 | P0C422, Q0D5P8, Q7FB12, A0A0P0Y0Q1, Q69SV0 |
| | Organelle outer membrane | GO:0031968 | 2.55E-03 | 25 | 5 | P0C422, Q0D5P8, Q7FB12, A0A0P0Y0Q1, Q69SV0 |
| | Outer membrane | GO:0019867 | 2.60E-03 | 25 | 5 | P0C422, Q0D5P8, Q7FB12, A0A0P0Y0Q1, Q69SV0 |
| | Plastid membrane | GO:0042170 | 5.76E-03 | 25 | 5 | P0C422, Q0D5P8, Q7FB12, A0A0P0Y0Q1, Q69SV0 |
| | Plastid envelope | GO:0009526 | 1.21E-02 | 25 | 5 | P0C422, Q0D5P8, Q7FB12, A0A0P0Y0Q1, Q69SV0 |
| | Photosystem II | GO:0009523 | 3.08E-04 | 25 | 4 | P0C422, Q6EP57, Q0D5P8, Q6Z8N7 |
| | Photosystem | GO:0009521 | 6.26E-04 | 25 | 4 | P0C422, Q6EP57, Q0D5P8, Q6Z8N7 |
| | Photosystem II oxygen evolving complex | GO:0009654 | 8.69E-04 | 25 | 3 | Q6EP57, Q0D5P8, Q6Z8N7 |
| | Extrinsic component of membrane | GO:0019898 | 1.47E-02 | 25 | 3 | Q6EP57, Q0D5P8, Q6Z8N7 |
| | Oxidoreductase complex | GO:1990204 | 1.66E-02 | 25 | 3 | Q6EP57, Q0D5P8, Q6Z8N7 |
| KEGG | Photosynthesis | KEGG:00195 | 8.35E-06 | 13 | 5 | P0C422, A0A0P0W0Z6, Q6EP57, Q0J8M2, Q0D5P8 |

Enrichment categories of exclusively downregulated proteins ($\log_2$ FC < 1; q-value<0.05). Abbreviations: GO, Gene Ontology; MF, Molecular Function; BC, Biological Process; *Padj, p*-value adjusted for multiple testing using the g:SCS algorithm (*Padj* < 0.05); Q, number of upregulated proteins into Source category; T∩Q, number of overlapping proteins within the specific Term ID.
